# Proteolytic control of mitochondrial calcium transport by intermembrane-space proteases

**DOI:** 10.64898/2026.08.20.745762

**Authors:** Anupriya Sinha, Kunal Samantaray, Ashlesha Kadam, Pooja Jadiya, Dhanendra Tomar

**Affiliations:** Department of Cardiovascular Medicine, Wake Forest University School of Medicine, Winston-Salem, NC 27157 USA; Department of Internal Medicine, Section of Gerontology and Geriatric Medicine, Wake Forest University School of Medicine, Winston-Salem, NC 27157 USA

**Keywords:** Mitochondria, Calcium, mtCU, MCU, MICU, NCLX, Proteases

## Abstract

The mitochondrial intermembrane space (IMS) is a critical regulatory interface for mitochondrial calcium (_m_Ca^2+^) flux. Positioned between the outer and inner mitochondrial membranes, the IMS links cytosolic Ca^2+^ signal to regulated Ca^2+^ uptake into the matrix. This positioning allows the IMS to influence _m_Ca^2+^ transport and Ca^2+^-dependent mitochondrial metabolism. _m_Ca^2+^ homeostasis is governed mainly by the mitochondrial calcium uniporter complex (mtCU), which mediates _m_Ca^2+^ uptake, and the Na^+^/Ca^2+^ exchanger NCLX, which drives _m_Ca^2+^ efflux. However, whether IMS regulatory events, particularly proteolytic remodeling by IMS proteases, control this transport machinery remains unclear. Using complementary knockout and overexpression approaches targeting ten IMS proteases (NLN, ATP23, IMMP1L, IMMP2L, YME1L1, OMA1, LACTB, LACTB2, PARL, and HTRA2), we identified protease-specific remodeling of mtCU components and NCLX abundance. Transcriptomic and proteomic analyses showed that these changes arise largely from protease-specific control of transporter stability rather than transcriptional regulation alone. Proximity-labeling proteomics further revealed spatial associations between IMS proteases and _m_Ca^2+^ transport components. Functionally, perturbing IMS proteases altered _m_Ca^2+^ flux and reduced _m_Ca^2+^ retention capacity, indicating impaired buffering against Ca^2+^ overload. Together, these findings identify IMS proteases as a proteostatic regulatory network controlling _m_Ca^2+^ transport and establish a mechanistic link between mitochondrial proteostasis and Ca^2+^ homeostasis.

Graphical abstract
IMS proteases regulate mitochondrial calcium uniporter complex architecture and mitochondrial Ca²⁺ homeostasis.
Systematic perturbation of intermembrane space (IMS) proteases by knockout (KO) and overexpression (OE) reveals a regulatory network linking IMS proteostasis to the mitochondrial calcium uniporter (mtCU) machinery. Changes in mtCU components were evaluated using complementary approaches, including mtCU protein expression analysis, transcriptomics, cycloheximide-based degradation proteomics to assess protein stability and turnover, and UltraID-based proximity proteomics to identify potential IMS protease-mtCU interactions. Integration of these datasets identifies multiple IMS proteases as regulators of the expression, stability, and organization of mtCU components. In the network schematic, the “+” symbol indicates positive regulation of an mtCU protein by the indicated IMS protease, whereas the “−” symbol indicates negative regulation. Perturbation of IMS protease expression consequently remodels the mtCU machinery, disrupts _m_Ca^2+^ uptake and efflux balance, impairs _m_Ca^2+^ buffering capacity, and increases susceptibility to _m_Ca^2+^ overload and mitochondrial permeability transition pore (mPTP) opening. Together, these findings establish IMS proteases as an integrated proteostatic network that maintains mtCU complex architecture and mitochondrial Ca^2+^ homeostasis.

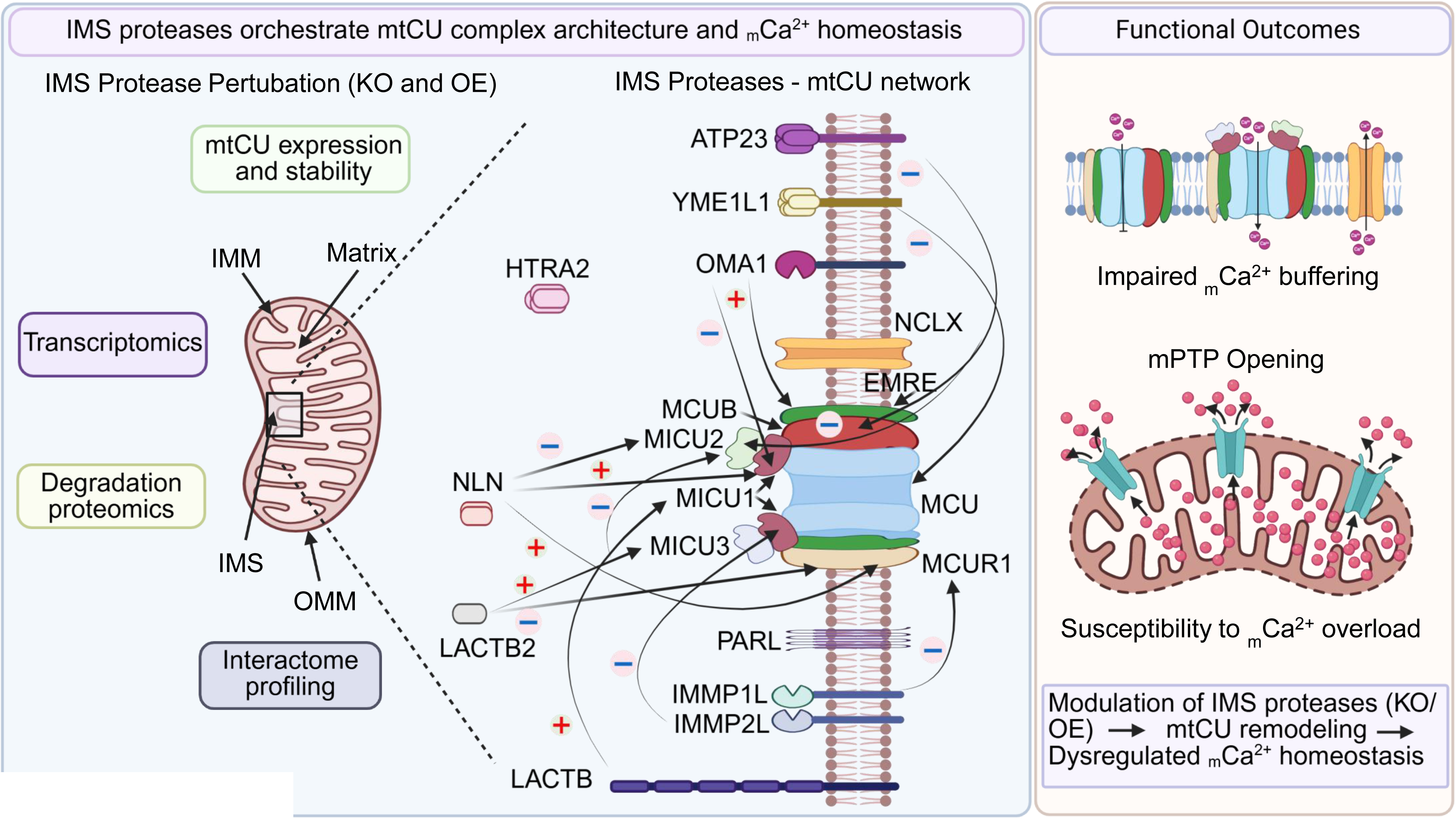

## Introduction

Calcium (Ca^2+^) is a versatile intracellular second messenger that coordinates diverse cellular processes, including metabolism, gene expression, secretion, excitability, and cell fate decisions^1–7^. Maintaining Ca^2+^ homeostasis is therefore critical for enabling cells to adapt to changing metabolic demands^8–12^. Mitochondria play a central role in decoding intracellular Ca^2+^ signals. Rather than serving merely as passive Ca^2+^ buffers, mitochondria actively regulate Ca^2+^ uptake and release, shaping cytosolic Ca^2+^ transients and coupling Ca^2+^ signaling to cellular bioenergetics^9,10,12–15^. Within the mitochondrial matrix, Ca^2+^ activates key dehydrogenases of the tricarboxylic acid cycle, enhances oxidative phosphorylation, and supports ATP production, allowing cells to rapidly adjust to energetic demand^9,10,12–15^. Conversely, excessive mitochondrial Ca^2+^ (_m_Ca^2+^) accumulation promotes reactive oxygen species production, opening of the mitochondrial permeability transition pore, and activation of cell-death pathways^9,10,12,13^. Thus, mitochondrial Ca^2+^ transport must be tightly controlled to balance metabolic adaptation with protection from Ca^2+^ overload.

For Ca²⁺ to reach the mitochondrial matrix, it must cross two membranes with markedly different permeability properties. The outer mitochondrial membrane (OMM) is relatively permeable to small ions and metabolites because it contains porin-like voltage-dependent anion channels (VDAC1–3), which permit Ca^2+^ movement from the cytosol into the intermembrane space (IMS)^10,12,16^. In contrast, the inner mitochondrial membrane (IMM) is highly impermeable to ions, including Ca^2+^, owing to its protein-rich, tightly sealed lipid bilayer and the need to preserve the electrochemical proton gradient that drives oxidative phosphorylation. Consequently, Ca^2+^ entry across the IMM is tightly regulated and represents the rate-limiting step of _m_Ca^2+^ uptake, mediated predominantly by the mitochondrial calcium uniporter complex (mtCU)^10,12,16–20^. Together, these membrane properties make the IMS a key regulatory interface where Ca^2+^ crossing the permissive OMM is sensed and gated before entering the matrix through the selective IMM.

The mtCU is a multi-protein complex that mediates Ca^2+^ uptake from the IMS into the mitochondrial matrix^12,17–19^. Its pore-forming subunit, mitochondrial calcium uniporter (MCU), forms the central Ca^2+^-conducting channel^17–19^, whereas MCUB acts as a dominant-negative paralog that limits Ca^2+^ conductance^12,21,22^. Essential MCU regulator (EMRE) stabilizes MCU and functionally couples the pore to its regulatory subunits^23–25^, while mitochondrial calcium uniporter regulator 1 (MCUR1) supports mtCU assembly and optimal channel activity^26–28^. The mitochondrial calcium uptake (MICU) proteins MICU1, MICU2, and MICU3, are positioned on the IMS-facing side of the IMM and act as Ca^2+^-sensitive gatekeepers^29–43^. At resting Ca^2+^ levels, MICU1 keeps the channel closed, preventing improper Ca^2+^ entry. When cytosolic Ca^2+^ rises, this inhibition is relieved, and MICU1 together with MICU2 promotes rapid Ca^2+^ uptake. MICU3, enriched in excitable tissues, enhances cooperative Ca^2+^ uptake through MICU1-containing regulatory complexes. Matrix Ca^2+^ accumulation is counterbalanced by efflux mechanisms, primarily mediated by the mitochondrial Na^+^/Ca^2+^ exchanger NCLX, which extrudes Ca^2+^ from the matrix to maintain homeostasis^44–46^. Thus, the dynamic balance between mtCU-mediated influx and NCLX-mediated efflux determines net _m_Ca^2+^ levels and strongly influences mitochondrial metabolism, stress responses, and disease-associated mitochondrial dysfunction.

Importantly, many regulatory components of mtCU either reside within the IMS or are embedded in the IMM with domains exposed to the IMS^12,20^. The IMS is also enriched with a diverse repertoire of mitochondrial proteases that control protein maturation, mitochondrial quality control (MQC), and the degradation of misfolded or unassembled proteins^47–52^. Key IMS and IMM-associated proteases include yeast mtDNA escape 1-like (YME1L1)^53–56^, overlapping activity with m-AAA protease 1 (OMA1)^57^, lactamase beta and lactamase beta 2 (LACTB and LACTB2)^58,59^, presenilin-associated rhomboid-like (PARL)^60^, high-temperature requirement A2 (HTRA2)^61^, neurolysin (NLN)^62^, ATP23^63^, and inner mitochondrial membrane peptidase subunits 1 and 2 (IMMP1L and IMMP2L)^64^. This close spatial overlap raises the possibility that IMS proteases directly or indirectly regulate the stability, processing, turnover, or assembly of mtCU components. Thus, proteolytic control within the IMS may represent an important but underexplored mechanism for fine-tuning mtCU activity and _m_Ca^2+^ homeostasis.

Consistent with this idea, altered expression of the IMS protease PARL has been shown to significantly affect _m_Ca^2+^ uptake^65^. Although PARL does not appear to directly interact with MICU1 or MICU2, PARL modulation changes the monomeric and dimeric forms of these proteins, suggesting indirect regulation of mtCU components. In addition, YME1L1 has been reported to degrade MICU1 and MICU2 when they are not properly assembled within mtCU, highlighting an MQC mechanism that links IMS proteolysis to mtCU regulation^56^. These studies suggest that IMS proteases may influence _m_Ca^2+^ signaling by regulating the abundance and assembly state of mtCU components. However, whether IMS proteases act broadly as a coordinated regulatory network controlling mtCU and NCLX expression, stability, and function remains unclear.

Here, we systematically investigated how IMS proteases regulate the _m_Ca^2+^ transport machinery. Using complementary knockout and overexpression approaches targeting ten IMS proteases, we assessed mtCU components and NCLX through protein-expression analysis, RNA sequencing, cycloheximide(CHX)-based proteostasis profiling, UltraID proximity proteomics, and functional measurements of _m_Ca^2+^ uptake, efflux, and calcium retention capacity. Our findings identify IMS proteases as an interconnected proteostatic regulatory network that shapes the abundance, stability, spatial organization, and activity of the _m_Ca^2+^ transport machinery. These results provide a mechanistic framework linking mitochondrial protein quality control within the IMS to _m_Ca^2+^ homeostasis.

## Results

### IMS proteases remodel mtCU and NCLX protein abundance

To determine whether IMS proteases regulate the _m_Ca^2+^ transport machinery, we generated HEK293T knockout (KO) cell lines lacking individual IMS proteases, including NLN, ATP23, IMMP1L, IMMP2L, YME1L1, OMA1, LACTB, LACTB2, PARL, or HTRA2. Complementary overexpression studies were performed using Flag-tagged IMS proteases. Successful KO and OE of the targeted IMS proteases was confirmed by immunoblotting (**Supplementary Fig. 1A, B**). We then examined the abundance of mtCU components, including MCU, MCUB, MICU1, MICU2, MICU3, MCUR1, and EMRE, together with NCLX. Loss of individual IMS proteases caused distinct changes in the abundance of multiple mtCU components and NCLX (**Fig. 1A**), which were quantified by densitometry and summarized as a heatmap (**Fig. 1B**). Statistical analyses for each mtCU component across the KO models are provided in **Supplementary Table 1**. Similar protease-specific remodeling was observed after IMS protease OE (**Fig. 1C, D**), indicating that knockout and overexpression perturbations dynamically alter the composition of the _m_Ca^2+^ transport machinery. Statistical analyses for the OE datasets are summarized in **Supplementary Table 2**. To integrate the KO and OE datasets, we generated a radar plot comparing the effects of each IMS protease perturbation on mtCU and NCLX expression (**Fig. 1E**). *NLN* deficiency markedly reduced nearly all mtCU components while increasing the inhibitory subunits MCUB and MICU2. In contrast, deletion of *IMMP2L* or *OMA1* broadly increased the expression of mtCU components. *YME1L1* and *LACTB* KO similarly increased most mtCU proteins, except MICU3 in *YME1L1* KO cells and MICU1/MICU3 in *LACTB* KO cells. Conversely, ATP23 OE reduced all measured mtCU components, whereas manipulation of the remaining IMS proteases produced component-specific effects on mtCU protein abundance.

**Figure 1.**
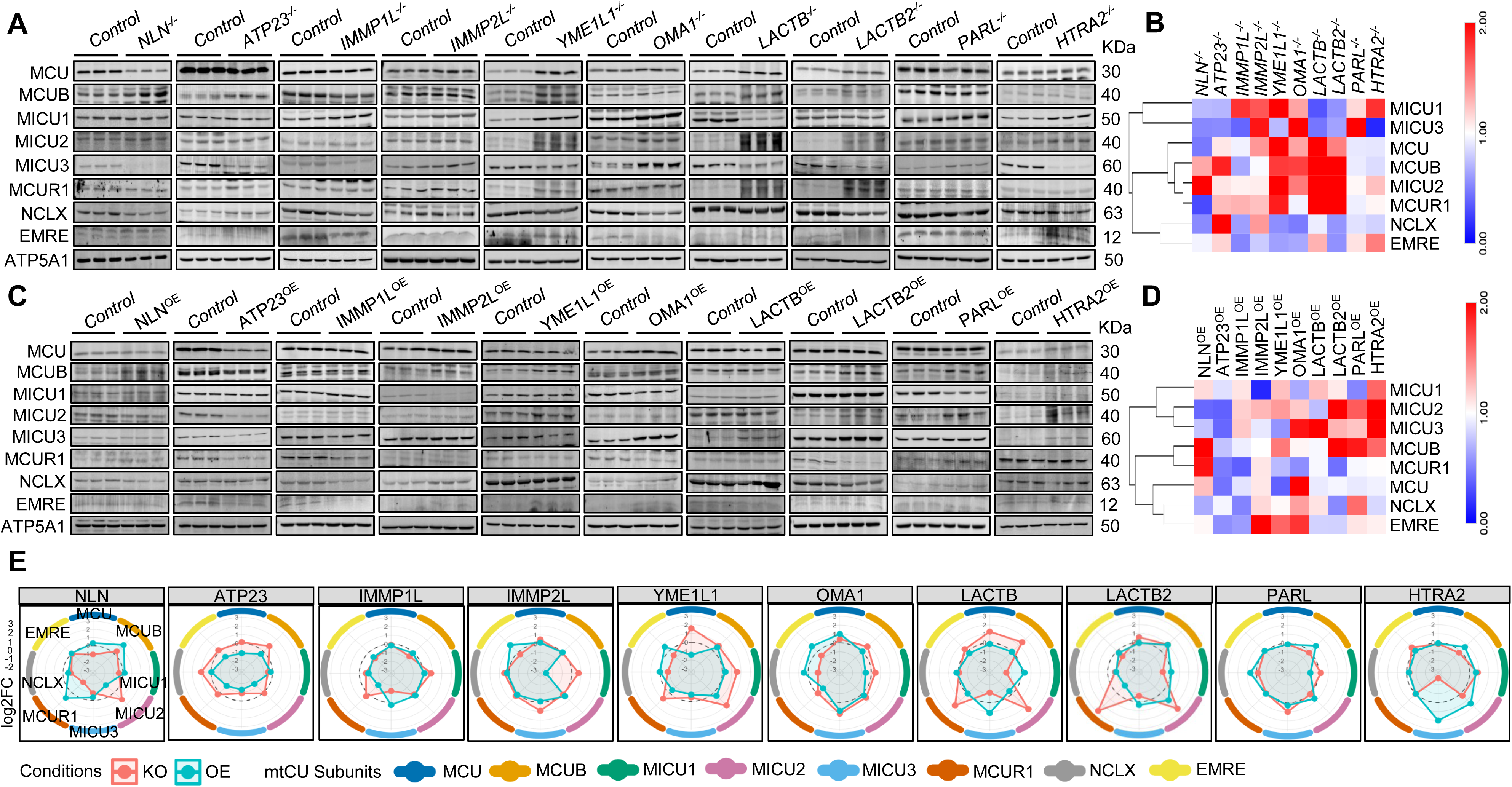
IMS proteases remodel the _m_Ca^2+^ transport machinery. **A.** Representative western blots showing the abundance of mtCU proteins MCU, MCUB, MICU1, MICU2, MICU3, MCUR1, and EMRE, together with NCLX, in HEK293T control cells and IMS protease KOs lacking *NLN*, *ATP23*, *IMMP1L*, *IMMP2L*, *YME1L1*, *OMA1*, *LACTB*, *LACTB2*, *PARL*, and *HTRA2*. **B.** Heatmap showing densitometric quantification of mtCU/NCLX abundance in IMS protease KO cells. Protein levels were quantified from western blots using ImageJ, normalized to the corresponding loading control, and expressed relative to HEK293T control cells. **C.** Representative western blots showing mtCU/NCLX expression following OE of Flag-tagged IMS proteases in HEK293T cells. **D.** Heatmap showing ImageJ-based densitometric quantification of mtCU components and NCLX abundance in IMS protease OE models, normalized to loading control and expressed relative to vector-transfected control cells. **E.** Spider radar plot integrating KO and OE datasets to compare protease-dependent changes in mtCU and NCLX expression. Values are shown as log_2_ fold change (log_2_FC) relative to the corresponding control condition, highlighting reciprocal and condition-specific regulation of _m_Ca^2+^ transport proteins by individual IMS proteases. Quantification was performed from three independent biological replicates.

A complementary Sankey network further revealed extensive connectivity between IMS proteases and individual mtCU components, with each protease displaying a distinct regulatory signature (**Supplementary Fig. 1C**). This analysis showed that multiple IMS proteases converge on common Ca^2+^ transport proteins, while individual proteases simultaneously influence several mtCU components. To classify these relationships, mtCU proteins were grouped into two major regulatory classes (**Supplementary Fig. 1D**): reciprocal positive regulators, whose expression decreased after KO and increased after OE, and reciprocal negative regulators, whose expression showed the opposite pattern. Correlation analyses further demonstrated coordinated regulation among mtCU proteins in both KO and OE models (**Supplementary Fig. 1E, F**). Collectively, these findings establish IMS proteases as an interconnected regulatory network that differentially remodels the abundance of the _m_Ca^2+^ transport machinery.

### IMS proteases remodel mtCU proteostasis with limited transcriptional regulation and prominent proteostatic control

Having established that IMS protease perturbation remodels mtCU and NCLX protein abundance, we next analyzed RNA-seq datasets from all IMS protease KO cell lines to determine whether these changes were driven by transcriptional regulation or altered protein stability. RNA-seq heatmap analysis revealed only modest changes in transcript levels of mtCU components and NCLX (**Fig. 2A**). Examination of the corresponding adjusted *p-values* further indicated that most transcript changes were not statistically significant (**Supplementary Table 3**), suggesting that loss of individual IMS proteases has a limited effect on mtCU and NCLX gene expression. Correlation analysis revealed distinct co-expression relationships among mtCU components (**Fig. 2B**). MCU positively correlated with MICU1 and MCUB but negatively correlated with MICU2, whereas EMRE positively correlated with MICU2 and MICU3. These relationships suggest coordinated regulation of specific mtCU subunits despite the limited overall transcriptional response to IMS protease deletion. Consistent with this interpretation, protein-to-RNA ratio analysis revealed substantial differences among mtCU components across KO models (**Fig. 2C**), indicating that transcriptional changes alone do not fully explain the observed remodeling of the _m_Ca^2+^ transport machinery, pointing instead to a major contribution of protein stability.

**Figure 2.**
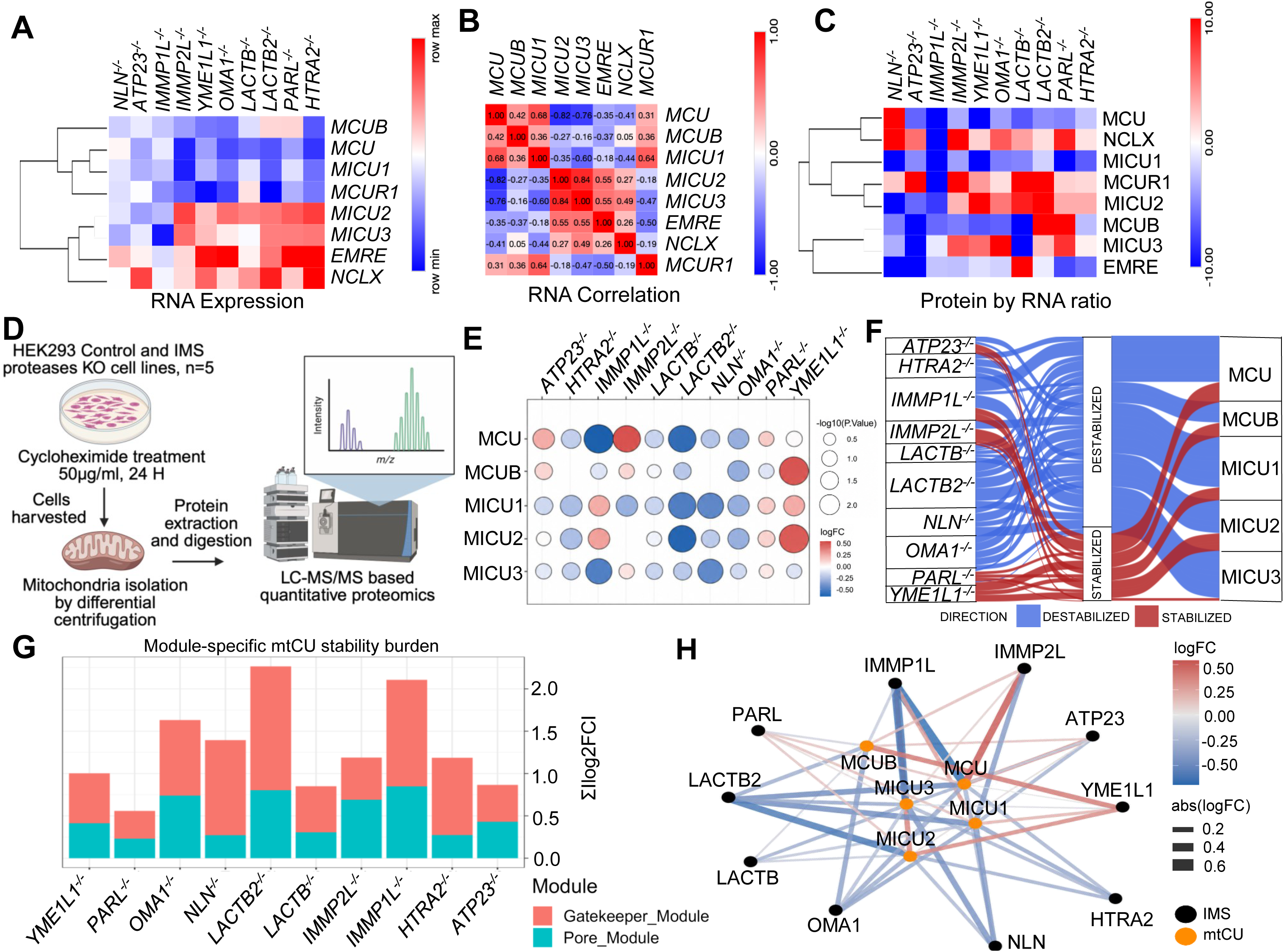
IMS proteases remodel mtCU/NCLX abundance through limited transcriptional changes and prominent proteostatic regulation. **A.** Heatmap showing RNA-seq-based expression changes of _m_Ca^2+^ transport genes across KO models. **B.** Correlation analysis of mtCU and NCLX transcript levels across KO models. **C.** Protein-to-RNA ratio analysis comparing mtCU/NCLX protein abundance with corresponding transcript levels across IMS protease KO models. **D.** Experimental workflow for CHX-based mitochondrial proteostasis analysis. **E.** Bubble plot showing stability changes of detectable mtCU proteins across KO models following CHX treatment. **F.** Sankey alluvial plot identifying IMS protease-dependent stabilization or destabilization of mtCU and NCLX proteins. **G.** Functional-module analysis classifying mtCU proteins into pore-forming components, including MCU and MCUB, and regulatory/gatekeeper components, including MICU1, MICU2, MICU3, MCUR1, and EMRE. **H.** Network analysis of CHX proteomics data showing regulatory connectivity between IMS proteases and mtCU/NCLX proteins. RNA-seq and CHX proteomics data were generated from four and five independent technical replicates, respectively.

To assess mtCU proteostasis more directly, cells were treated with CHX to inhibit de novo protein synthesis and then subjected to quantitative mitochondrial proteomics (**Fig. 2D**). MCUR1, NCLX, and EMRE were not detected in the CHX-proteomics dataset and therefore could not be evaluated for protein stability. Analysis of the remaining mtCU components revealed distinct proteostatic effects across IMS protease KO models (**Fig. 2E**). The CHX-proteomics dataset further identified protease-specific relationships between IMS proteases and mtCU components (**Fig. 2F**). Loss of *HTRA2, LACTB, LACTB2, NLN,* and *OMA1* was associated with broad reductions in mtCU protein stability, suggesting that these proteases contribute to maintenance of mtCU stability. In contrast, deletion of other IMS proteases selectively stabilized specific mtCU components. MCU showed increased relative stability in *ATP23*-, *IMMP2L*-, and *PARL* KO cells, whereas MCUB was stabilized in *ATP23*-, *IMMP2L*-, and *YME1L1* KO cells. MICU1 and MICU2 showed increased stability in *IMMP1L*-, *PARL*-, and *YME1L1* KO cells, while MICU3 showed increased relative stability specifically in *IMMP2L-* and *PARL* KO cells. Classification of mtCU proteins into pore-forming components (MCU and MCUB) and regulatory/gatekeeper modules (MICU1, MICU2, and MICU3) further showed that IMS proteases differentially regulate these functional subgroups (**Fig. 2G**). Network analysis highlighted extensive connectivity between IMS proteases and mtCU components, identifying convergent regulatory interactions that collectively shape mtCU proteostasis (**Fig. 2H**). Consistent with these findings, principal component analysis showed that each IMS protease KO produced a distinct mtCU proteostasis landscape (**Supplementary Fig. 2A**), and directional analysis revealed unique stabilizing and destabilizing effects of individual proteases on specific mtCU proteins (**Supplementary Fig. 2B**). Together, these findings demonstrate that IMS proteases exert broad and component-specific control over mtCU protein turnover, with some proteases contributing to mtCU stability and others promoting degradation of distinct mtCU components.

### IMS protease proximity interactomes reveal associations with the _m_Ca^2+^ transport machinery

To determine whether IMS proteases occupy spatial proximity to mtCU components, we performed proximity-dependent biotin labeling using UltraID-tagged IMS proteases followed by quantitative mass spectrometry **(Fig. 3A)**^66^. This approach enabled identification of proteins located within the immediate molecular environment of each IMS protease. UltraID proteomics detected multiple mtCU components in proximity to individual IMS proteases (**Fig. 3B**). PARL and ATP23 did not show detectable enrichment of mtCU components under these experimental conditions, whereas the remaining IMS proteases showed selective associations with distinct mtCU subunits. MCU was identified in proximity to LACTB, LACTB2, and YME1L1, while MCUB was detected in proximity to IMMP2L, LACTB, and NLN. Among the regulatory gatekeeper subunits, MICU1 was enriched with HTRA2, IMMP1L, IMMP2L, LACTB, NLN, OMA1, and YME1L1, whereas MICU2 was detected with IMMP1L, IMMP2L, LACTB, NLN, OMA1, and YME1L1. MCUR1 was identified in proximity to IMMP1L and NLN. In contrast, MICU3, EMRE, and NCLX were not detected in any IMS protease proximity dataset under the conditions tested.

**Figure 3.**
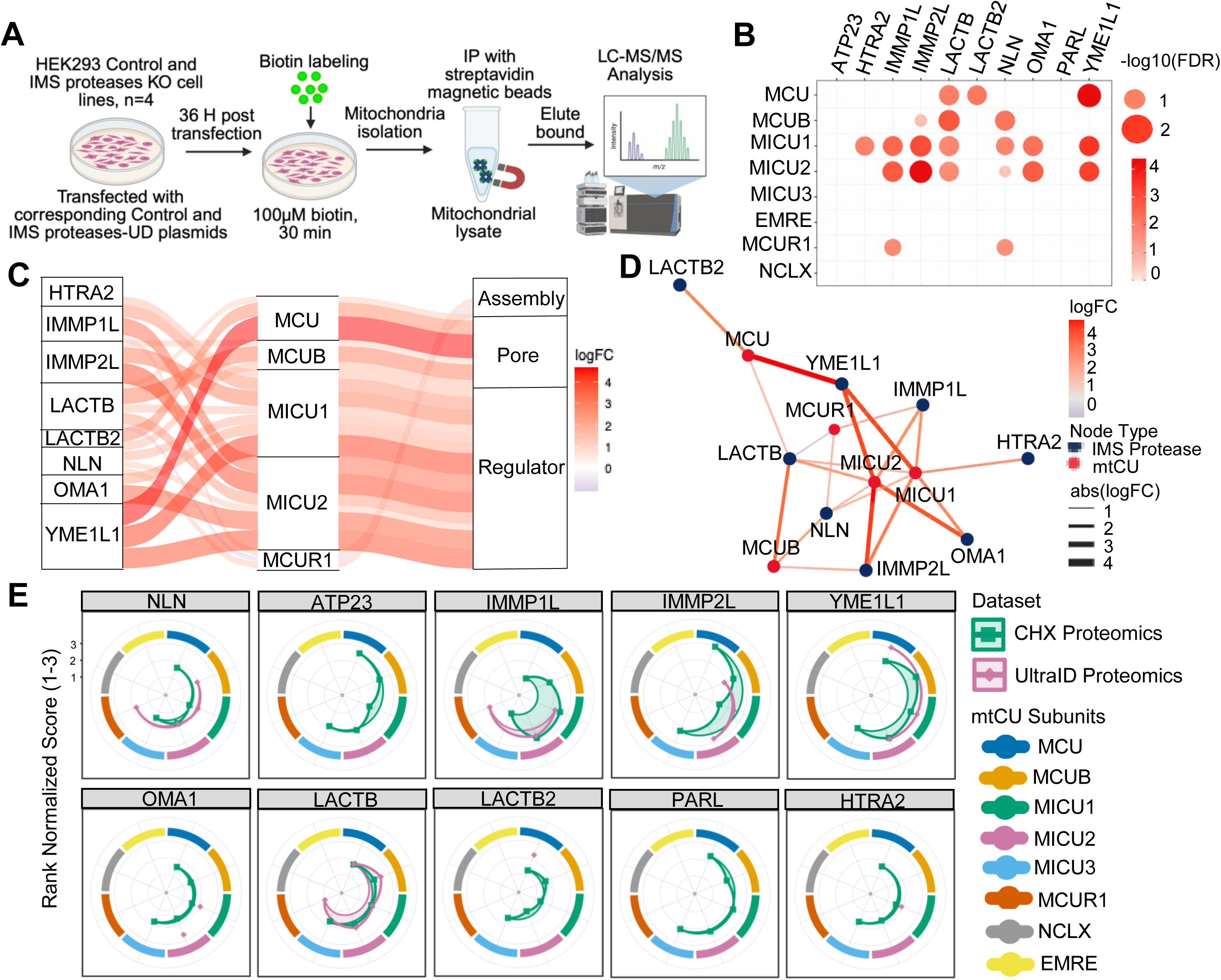
UltraID proximity proteomics links IMS proteases to the _m_Ca^2+^ transport machinery. **A.** Schematic workflow of the UltraID proximity-labeling proteomics experiment. **B.** Bubble plot showing UltraID-based proximity enrichment of mtCU/NCLX across individual IMS proteases. This analysis identifies protease-specific proximity relationships between IMS proteases and _m_Ca^2+^ transport proteins. **C.** Sankey alluvial plot visualizing IMS protease– mtCU/NCLX proximity relationships detected by UltraID proteomics. **D.** Network analysis of the UltraID proximity interactome showing connectivity between IMS proteases and mtCU/NCLX proteins. **E.** Spider radar plots integrating CHX-based proteostasis data and UltraID proximity proteomics for each IMS protease. Each radar plot summarizes the relationship between mtCU/NCLX protein stability and UltraID proximity enrichment across individual _m_Ca^2+^ transport components, revealing protease-specific coupling between physical proximity and post-translational regulation. UltraID data were generated from four independent biological replicates.

To visualize these relationships, we generated a Sankey network linking IMS proteases to their proximal mtCU proteins (**Fig. 3C**). This analysis showed that multiple IMS proteases converge on common mtCU components, particularly MICU1 and MICU2, while individual proteases associate with multiple mtCU subunits, revealing an interconnected proximity network. Network analysis further emphasized this connectivity and identified MICU1 and MICU2 as the most extensively connected hubs, followed by MCU and MCUB (**Fig. 3D**). We next integrated the UltraID proximity-proteomics dataset with CHX-based proteostasis data to determine whether spatial proximity between IMS proteases and mtCU components corresponded with changes in mtCU protein stability. Radar plots generated for each IMS protease revealed protease-specific relationships between proximity and proteostasis (**Fig. 3E**). Some IMS proteases were both positioned near mtCU components and influenced their stability, whereas others showed regulatory effects that may occur indirectly or through broader mitochondrial remodeling. Finally, to summarize the multidimensional regulatory landscape, we generated an integrated dot plot combining all four datasets: mtCU/NCLX protein-expression changes in IMS protease KO cells, protein-expression changes following IMS protease OE, CHX-based proteomics, and UltraID proximity proteomics (**Supplementary Fig. 3**). Together, these findings reveal a selective proximity landscape in which individual IMS proteases associate with distinct subsets of mtCU components, suggesting multiple protease-specific regulatory nodes within the _m_Ca^2+^ transport machinery.

### Loss of IMS proteases alters _m_Ca^2+^ flux dynamics

To determine whether IMS protease-dependent remodeling of the _m_Ca^2+^ transport machinery alters mitochondrial Ca^2+^ handling, we measured _m_Ca^2+^ flux in all IMS protease KO models using acute Ca^2+^ pulses. Under high-Ca^2+^ conditions, cells were challenged with 10 µM CaCl_2_, and representative _m_Ca^2+^ uptake and efflux traces were generated for each KO model (**Fig. 4A–J**). These traces revealed protease-specific changes in both uptake rates and the subsequent efflux phase, indicating that loss of individual IMS proteases differentially affects _m_Ca^2+^ transport dynamics. Quantification of the 10 µM CaCl_2_ flux assays showed that *NLN*, *ATP23*, *IMMP1L*, *YME1L1*, *OMA1*, *LACTB*, *PARL, and HTRA2* KO significantly reduced _m_Ca^2+^ uptake rates compared with control cells (**Fig. 4K**). Efflux analysis further showed that *NLN*, *ATP23*, *YME1L1*, *OMA1*, *LACTB*, and *PARL* KO significantly reduced _m_Ca^2+^ efflux (**Fig. 4L**). These findings indicate that multiple IMS proteases contribute to efficient _m_Ca^2+^ uptake and extrusion under high Ca^2+^ load.

**Figure 4.**
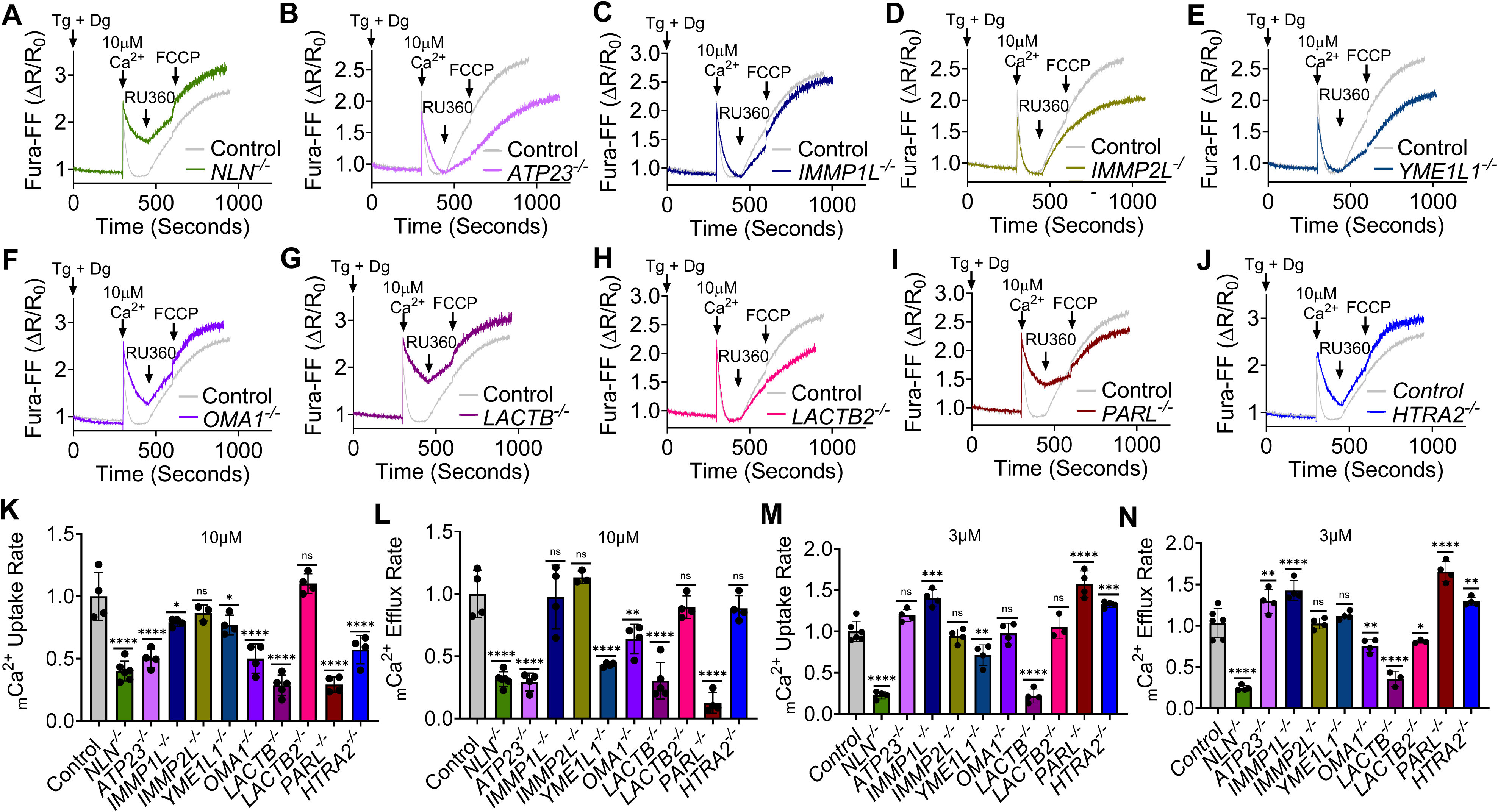
Loss of IMS proteases differentially alter _m_Ca^2+^ uptake and efflux dynamics. **A–J.** Representative _m_Ca^2+^ flux traces from HEK293T control cells and IMS protease KO cell lines following acute stimulation with 10 µM CaCl_2_. Traces show _m_Ca^2+^ uptake after CaCl_2_ addition and subsequent Ca^2+^ efflux dynamics in *NLN*, *ATP23*, *IMMP1L*, *IMMP2L*, *YME1L1*, *OMA1*, *LACTB*, *LACTB2*, *PARL*, and *HTRA2* KO cells compared with control cells. For comparison, the same representative control trace is displayed alongside each KO trace. **K.** Quantification of _m_Ca^2+^ uptake rates following 10 µM CaCl_2_ stimulation. **L.** Quantification of _m_Ca^2+^ efflux rates following 10 µM CaCl_2_ stimulation. **M.** Quantification of _m_Ca^2+^ uptake rates following low-calcium stimulation with 3 µM CaCl_2_. **N.** Quantification of _m_Ca^2+^ efflux rates following 3 µM CaCl_2_ stimulation. All flux quantifications were performed from a minimum of three independent biological replicates. Data are presented as mean ± SD. Statistical significance was determined using ordinary one-way ANOVA with multiple comparisons; *p < 0.05, **p < 0.01, ***p < 0.00, ****p < 0.0001, ns = not significant (p ≥ 0.05).

Because lower Ca^2+^ loads are more sensitive to MICU-dependent mtCU gating, we next measured _m_Ca^2+^ flux using 3 µM CaCl_2_. Representative uptake and efflux traces for each IMS protease KO model are shown in **Supplementary Fig. 4A–J**. Under these low-Ca^2+^ conditions, loss of IMS proteases again produced distinct effects on _m_Ca^2+^ flux. Quantification showed that *NLN*, *YME1L1*, and *LACTB* KO significantly reduced _m_Ca^2+^ uptake, whereas *IMMP1L*, *PARL*, and *HTRA2* KO increased uptake rates, indicating that IMS protease loss can either impair or enhance Ca^2+^ entry depending on the protease involved (**Fig. 4M**). Analysis of Ca^2+^ efflux at 3 µM CaCl_2_ showed reduced efflux in *NLN*, *OMA1*, *LACTB*, and *LACTB2* KO cells, while *ATP23*, *IMMP1L*, *PARL*, and *HTRA2* KO cells showed increased efflux (**Fig. 4N**). To integrate uptake and efflux behavior, we calculated uptake-to-efflux relationships under both calcium conditions. These analyses are shown for 10 µM CaCl_2_ and 3 µM CaCl_2_ in **Supplementary Fig. 4K** and **Supplementary Fig. 4L**, respectively. To provide a global comparison of Ca^2+^ transport phenotypes across all IMS protease KO models, we also generated integrated radar plots summarizing normalized _m_Ca^2+^ uptake and efflux rates at both 10 µM and 3 µM CaCl_2_ (**Supplementary Fig. 4M**). Collectively, these results demonstrate that IMS proteases are critical regulators of _m_Ca^2+^ flux dynamics. Loss of individual IMS proteases produces distinct effects on _m_Ca^2+^ uptake, efflux, and uptake-to-efflux coupling, indicating that IMS proteases regulate not only mtCU/NCLX abundance and stability but also the functional behavior of the _m_Ca^2+^ transport machinery.

### Overexpression of IMS proteases modulates _m_Ca^2+^ flux dynamics

To determine whether increased IMS protease abundance is sufficient to remodel _m_Ca^2+^ handling, we measured _m_Ca^2+^ uptake and efflux following OE of individual IMS proteases in HEK293T cells. Under high-Ca^2+^ conditions, cells were challenged with 10 µM CaCl_2_, and representative uptake and efflux traces were generated for each IMS protease OE model (**Fig. 5A–J**). These traces revealed protease-specific changes in _m_Ca^2+^ transport dynamics, indicating that elevated expression of individual IMS proteases is sufficient to modify both Ca^2+^ uptake and efflux. Quantification of the 10 µM CaCl_2_ flux assays showed that OE of YME1L1 and LACTB significantly reduced _m_Ca^2+^ uptake rates, whereas OE of NLN, ATP23, and IMMP1L increased uptake compared with control cells (**Fig. 5K**). Analysis of Ca^2+^ efflux under the same conditions further showed that YME1L1 and LACTB OE significantly reduced _m_Ca^2+^ efflux rates (**Fig. 5L**). These findings indicate that increased abundance of specific IMS proteases can either enhance or suppress _m_Ca^2+^ flux under high Ca^2+^ load. To determine whether IMS protease OE affects mtCU behavior under conditions more sensitive to MICU-dependent gating, we next measured _m_Ca^2+^ flux using a lower Ca^2+^ stimulus of 3 µM CaCl_2_. Representative uptake and efflux traces for each IMS protease OE model are shown in **Supplementary Fig. 5A–J**. Under these low-Ca^2+^ conditions, IMS protease OE again produced selective effects on _m_Ca^2+^ handling. Quantification showed that OE of IMMP1L, YME1L1, OMA1, and LACTB2 significantly reduced _m_Ca^2+^ uptake rates compared with control cells (**Fig. 5M**). Similarly, analysis of Ca^2+^ efflux at 3 µM CaCl_2_ demonstrated significantly reduced efflux following OE of ATP23, YME1L1, OMA1, and LACTB2 (**Fig. 5N**).

**Figure 5.**
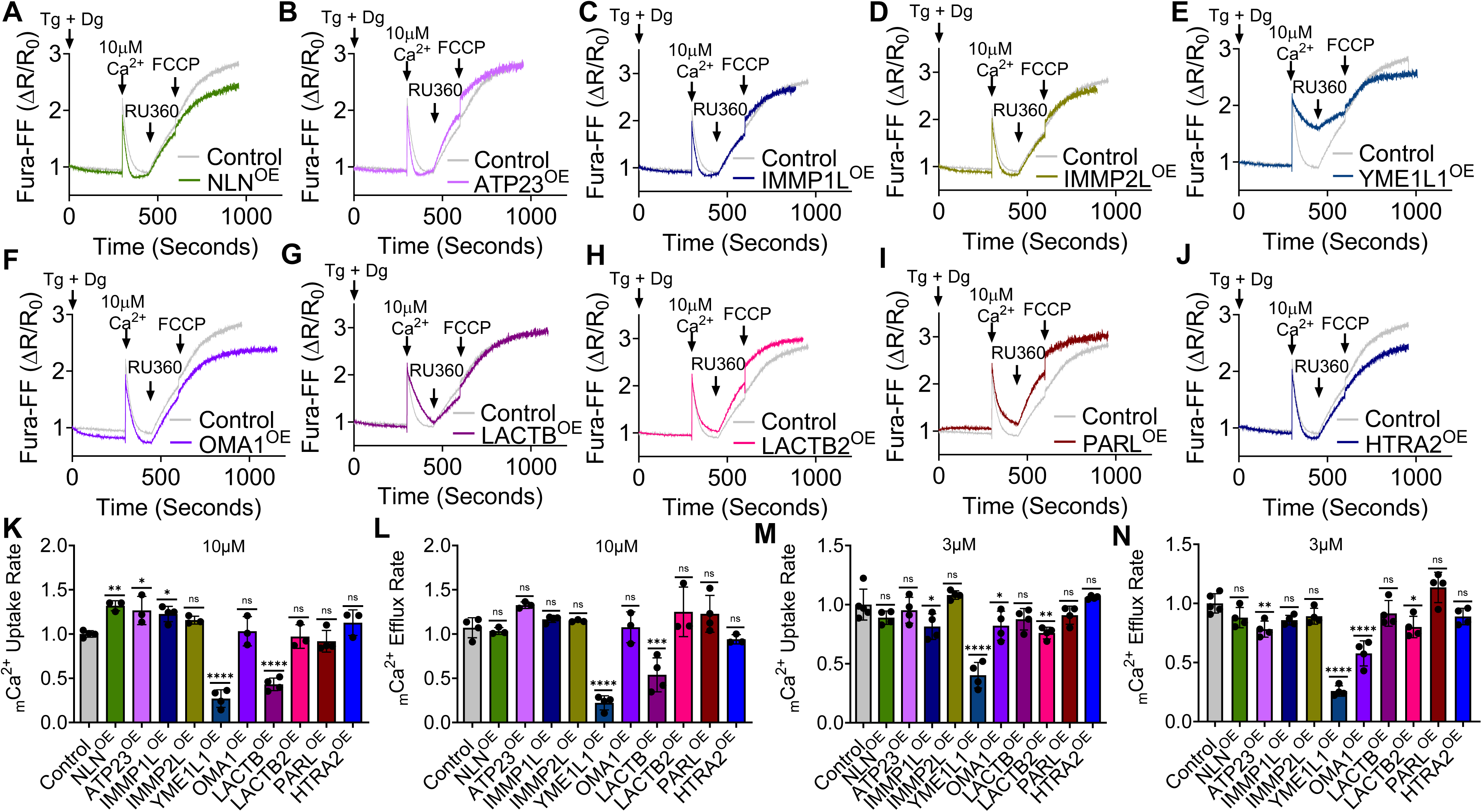
IMS protease overexpression differentially modulates _m_Ca^2+^ uptake and efflux dynamics. **A–J.** Representative _m_Ca^2+^ flux traces from HEK293T control cells and cells overexpressing individual IMS proteases following acute stimulation with 10 µM CaCl_2_. Traces show _m_Ca^2+^ uptake after CaCl_2_ addition and subsequent Ca^2+^ efflux dynamics in cells overexpressing NLN, ATP23, IMMP1L, IMMP2L, YME1L1, OMA1, LACTB, LACTB2, PARL, or HTRA2 compared with control cells. For comparison, the same representative control trace is displayed in all panels. **K.** Quantification of _m_Ca^2+^ uptake rates following 10 µM CaCl_2_ stimulation. **L.** Quantification of _m_Ca^2+^ efflux rates following 10 µM CaCl_2_ stimulation, demonstrating differential regulation of Ca^2+^ extrusion following IMS protease OE. **M.** Quantification of _m_Ca^2+^ uptake rates following low-calcium stimulation with 3 µM CaCl_2_. **N.** Quantification of _m_Ca^2+^ efflux rates following 3 µM CaCl_2_ stimulation, showing Ca^2+^-load-dependent effects of IMS protease OE on _m_Ca^2+^ extrusion. All flux quantifications were performed from a minimum of three independent biological replicates. Data are presented as mean ± SD. Statistical significance was determined using ordinary one-way ANOVA with multiple comparisons; *p < 0.05, **p < 0.01, ***p < 0.001, ****p < 0.0001, ns = not significant (p ≥ 0.05).

To integrate uptake and efflux behavior following IMS protease OE, we calculated uptake-to-efflux relationships under both Ca^2+^ conditions. These analyses are shown for 10 µM CaCl_2_ and 3 µM CaCl_2_ in **Supplementary Fig. 5K** and **Supplementary Fig. 5L**, respectively. An integrated radar plot comparing normalized _m_Ca^2+^ uptake and efflux rates across all IMS protease OE models under both 10 µM and 3 µM CaCl_2_ conditions is presented in **Supplementary Fig. 5M**. Together, these findings demonstrate that increasing IMS protease abundance is sufficient to reshape _m_Ca^2+^ uptake–efflux coupling, indicating that mitochondrial Ca²⁺ transport is sensitive to changes in IMS protease abundance.

### IMS proteases regulate mitochondrial Ca^2+^ retention capacity

Having shown that IMS protease loss alters _m_Ca^2+^ uptake and efflux, we next asked whether these changes affect mitochondrial Ca^2+^ retention capacity (CRC), a functional measure of mitochondrial Ca^2+^ buffering and resistance to Ca^2+^ overload. Representative CRC traces showed protease-specific changes in mitochondrial Ca^2+^ retention following repetitive Ca^2+^ pulses **(Fig. 6A-J)**. Quantitative analysis revealed that loss of *NLN*, *ATP23*, *IMMP1L*, *YME1L1*, *OMA1*, *LACTB*, *LACTB2*, *PARL*, and *HTRA2* significantly reduced CRC compared with control cells **(Fig. 6K)**, indicating a diminished capacity to retain Ca^2+^ during repetitive Ca^2+^ loading. In contrast, *IMMP2L* KO did not significantly alter CRC, suggesting that IMMP2L has a more limited or context-dependent role in regulating mitochondrial Ca^2+^ retention under these experimental conditions. Together, these findings indicate that most IMS proteases contribute to maintaining mitochondrial Ca^2+^ retention capacity.

**Figure 6.**
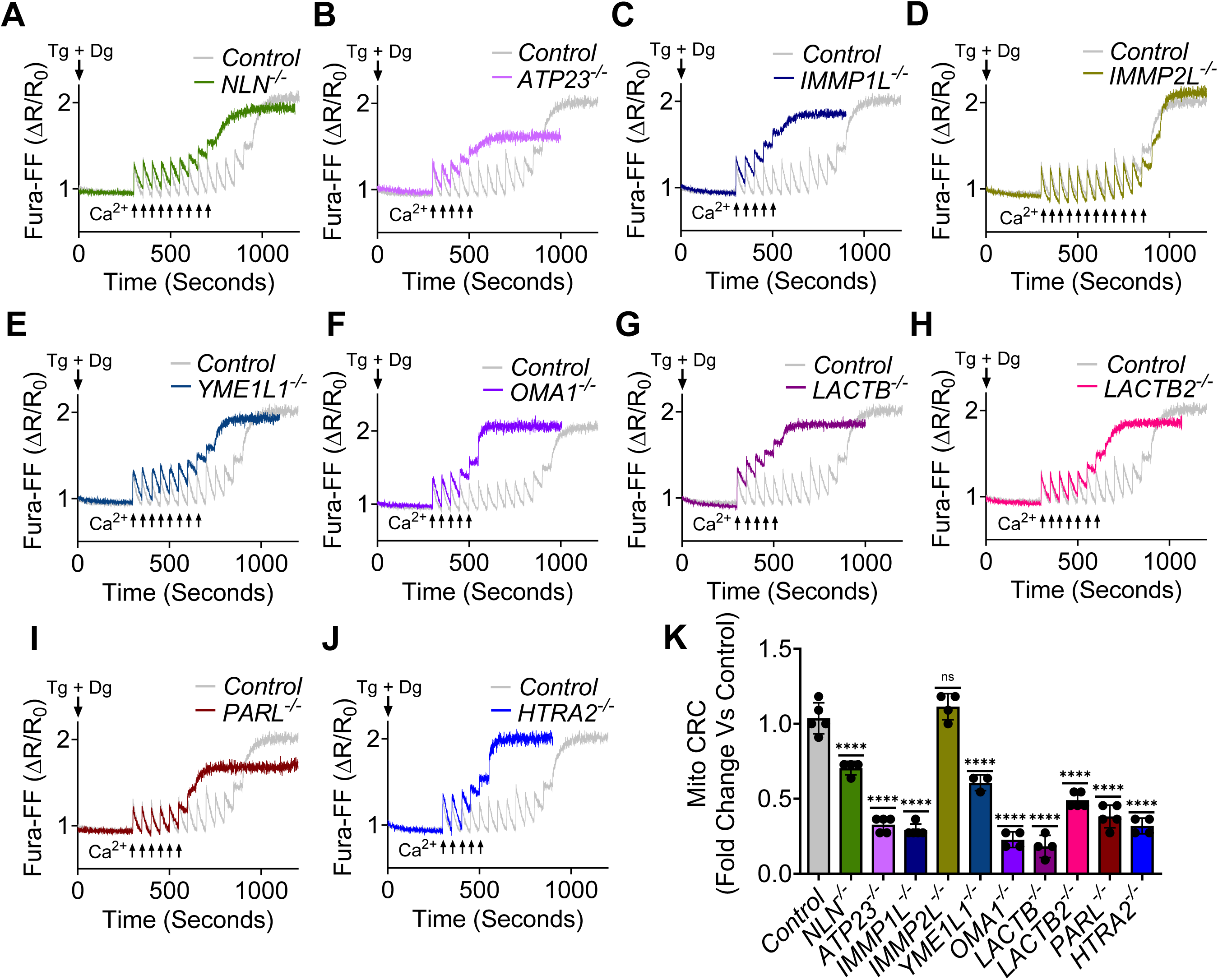
Loss of IMS proteases impairs mitochondrial CRC. **A–J.** Representative traces showing CRC responses in *NLN*, *ATP23*, *IMMP1L*, *IMMP2L*, *YME1L1*, *OMA1*, *LACTB*, *LACTB2*, *PARL*, and *HTRA2* KO cells compared with control cells. For comparison, the same representative control trace is displayed in panels B–E and G–J, whereas a second representative control trace is displayed in panels A and F. **K.** Quantification of mitochondrial CRC across IMS protease KO models. All quantitative analyses were performed using all independent control and KO biological replicate traces, including the representative control traces shown in panels A–J, and were not restricted to the representative traces presented in the figure. Quantification was performed from at least three independent biological replicates. Data are presented as mean ± SD. Statistical significance was determined using ordinary one-way ANOVA with multiple comparisons; ****p < 0.0001, ns = not significant (p ≥ 0.05).

Consistent with the flux phenotypes observed in Figures 4 and 5, impaired CRC following IMS protease deletion indicates that disruption of the IMS protease network compromises _m_Ca^2+^ homeostasis and increases susceptibility to Ca^2+^ overload. To determine whether increased IMS protease abundance also influences mitochondrial Ca^2+^ retention, we measured CRC following OE of individual IMS proteases. Representative CRC traces for all IMS protease OE models are shown in **Fig. 7A-J**. Quantitative analysis demonstrated that OE of NLN, ATP23, IMMP2L, YME1L1, OMA1, LACTB, LACTB2, PARL, and HTRA2 significantly reduced CRC compared with control cells (**Fig. 7K**), indicating a diminished capacity of mitochondria to retain Ca^2+^ during repetitive Ca^2+^ loading. In contrast, IMMP1L OE significantly increased CRC, suggesting enhanced mitochondrial Ca^2+^ buffering capacity. Together, these findings demonstrate that mitochondrial Ca^2+^ retention is sensitive to both loss and increased abundance of IMS proteases.

**Figure 7.**
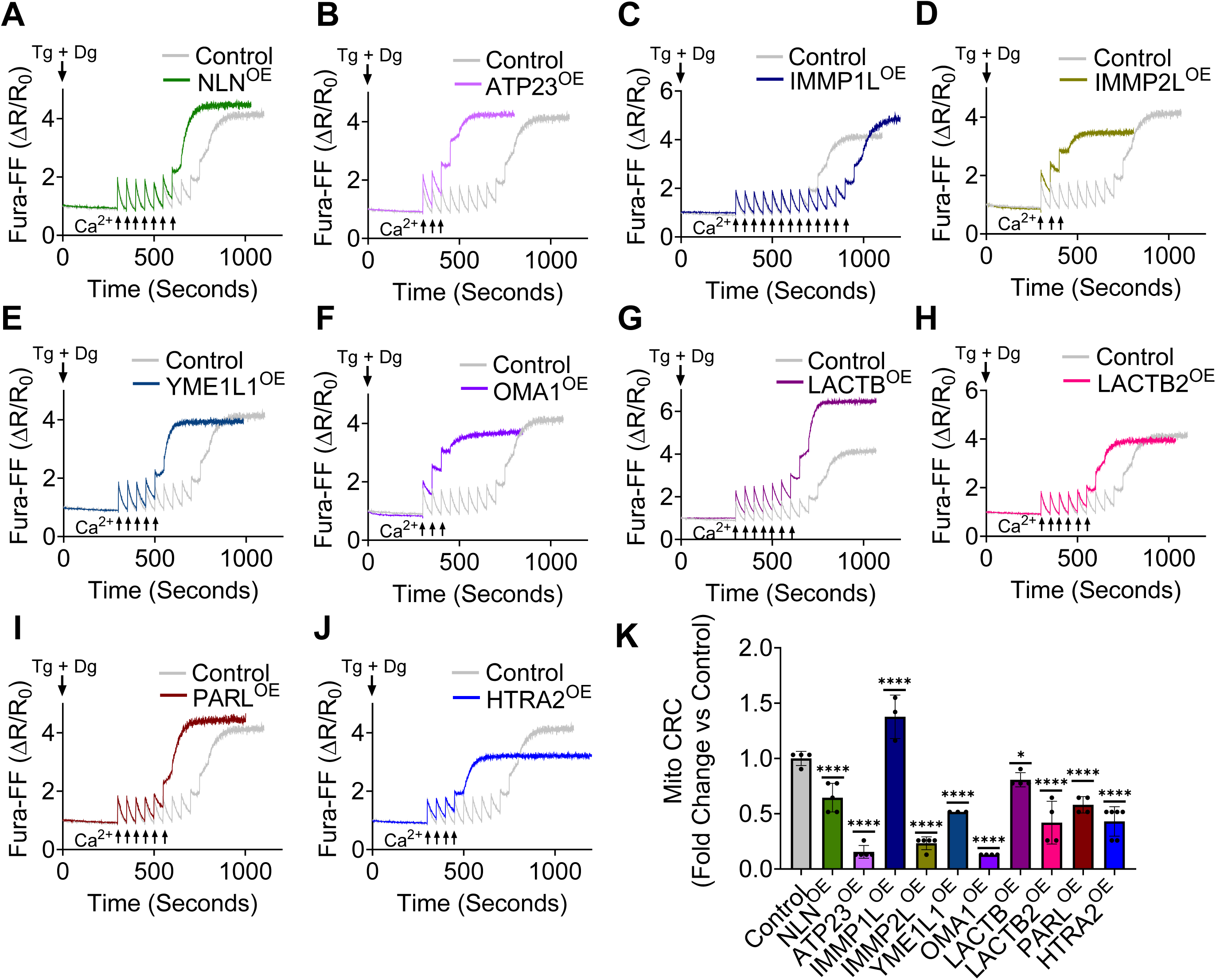
IMS protease OE alters mitochondrial CRC. **A–J.** Representative traces showing CRC responses in cells overexpressing NLN, ATP23, IMMP1L, IMMP2L, YME1L1, OMA1, LACTB, LACTB2, PARL, or HTRA2 compared with control cells. For comparison, the same representative control trace is displayed in all panels. **K.** Quantification of mitochondrial CRC following IMS protease OE. All quantitative analyses were performed using all independent control and IMS protease OE biological replicate traces and were not restricted to the representative traces shown in panels A–J. Quantification was performed from at least three independent biological replicates. Data are presented as mean ± SD. Statistical significance was determined using ordinary one-way ANOVA with multiple comparisons; *p < 0.05, ****p < 0.0001.

## Discussion

This study identifies IMS proteases as previously unrecognized regulators of the _m_Ca^2+^ transport machinery. By integrating transcriptomic, proteomic, proximity-labeling, and functional approaches, we show that IMS proteases control the abundance, stability, and activity of mtCU components and NCLX, thereby reshaping _m_Ca^2+^ uptake, efflux, and CRC. Although individual IMS proteases such as PARL have been linked to MICU1 and MICU2 regulation^65^, and stress-responsive proteases such as OMA1 and YME1L1 are established modulators of mitochondrial proteostasis^48,49,52,56,67–70^, the broader contribution of IMS proteases to _m_Ca^2+^ transport has remained unclear. Our findings extend this framework by defining IMS proteases as an interconnected regulatory network that couples mitochondrial proteostasis to Ca^2+^ signaling.

IMS proteases did not act through a single shared mechanism. Instead, each protease produced distinct effects on mtCU protein abundance and stability, supporting protease-specific regulatory programs that shape _m_Ca^2+^ uptake and efflux. The divergent phenotypes observed after protease loss and overexpression reveal a network of positive and negative relationships that shape mtCU abundance and likely influence complex organization. RNA-seq detected protease-specific changes in mtCU and NCLX transcripts, but these changes did not fully account for protein remodeling, indicating that post-transcriptional proteostatic regulation is a major component of this control. CHX-chase proteomics confirmed that IMS proteases regulate mtCU protein turnover, establishing protein stability as a key mechanism underlying remodeling of the _m_Ca^2+^ transport machinery. Several steady-state changes observed by western blotting were reproduced in the proteomics dataset, including accumulation of MCUB, MICU1, and MICU2 in *YME1L1* KO cells; reduced MCU, MICU1, and MICU3 in *NLN* KO cells; increased MCU and MCUB in *ATP23* deficiency; accumulation of MICU1 and MICU2 in *IMMP1L* deficiency; increased MCU in *IMMP2L* KO cells; and accumulation of MICU1 and MICU3 in *PARL* KO cells. This concordance supports post-translational regulation as a central determinant of mtCU abundance. At the same time, several MICU proteins were reduced in CHX-chase proteomics relative to steady-state measurements, likely reflecting their dynamic turnover and dependence on assembly for stability. Properly assembled MICU complexes are protected from proteolysis, whereas unassembled MICU proteins are susceptible to degradation by IMS proteases such as YME1L1^56^. Thus, the proteolytic sensitivity of MICU1, MICU2, and MICU3 may explain their greater depletion during CHX treatment and highlight mitochondrial proteostasis as a critical regulator of mtCU abundance, organization, and _m_Ca^2+^ homeostasis.

The proximity-labeling dataset provides systematic evidence that multiple IMS proteases reside near components of the _m_Ca^2+^ transport machinery, supporting localized protease-dependent regulation. PARL and ATP23 did not show detectable mtCU proximity partners under these experimental conditions, suggesting that their KO and OE effects may occur indirectly through maturation pathways or secondary proteostatic networks. In contrast, MICU1 and MICU2 were identified in proximity datasets for nearly all IMS proteases, placing these IMS-facing Ca^2+^-sensing gatekeepers at a key interface between mitochondrial proteostasis and Ca^2+^ signaling. The functional data show that IMS proteases regulate not only mtCU and NCLX abundance but also _m_Ca^2+^ transport activity. Under high Ca^2+^ stimulation (10 µM CaCl_2_), loss of most IMS proteases reduced _m_Ca^2+^ uptake, with the exception of *IMMP2L* and *LACTB2* KO cells. Preserved uptake in these models may reflect compensatory increases in MCU and MCUR1; in *IMMP2L* KO cells, elevated MICU3 may further support _m_Ca^2+^ influx.

Because high Ca^2+^ stimulation may primarily report maximal channel activity and mask changes in the activation threshold^30,31,35,38^, we also assessed flux at 3 µM CaCl_2_, a concentration closer to the MICU-dependent activation range of mtCU. MICU1 maintains channel closure at resting Ca^2+^ and promotes opening upon Ca^2+^ binding, MICU2 reinforces inhibition at low Ca^2+^, and MICU3 enhances _m_Ca^2+^ uptake^30,31,33,35,38–41,43^. Under low-Ca^2+^ conditions, _m_Ca^2+^ uptake was reduced in *NLN*, *YME1L1*, and *LACTB* KO cells but enhanced in *IMMP1L*, *PARL*, and *HTRA2* KO cells. Reduced uptake in *NLN*, *LACTB*, and *YME1L1* KO cells is consistent with increased inhibitory gatekeeping driven by elevated MICU2 and/or reduced MICU3. In contrast, increased uptake in *IMMP1L* and *HTRA2* KO cells may reflect selective MICU1 accumulation, which could favor MICU1-MICU1 homodimers that activate more readily than MICU1-MICU2 heterodimers^35,38,40,71^. Similarly, increased MICU3 in *PARL* KO cells may enhance _m_Ca^2+^ entry through MICU1-MICU3 regulatory heterodimers.

OE studies further demonstrated that increased IMS protease abundance is sufficient to remodel mtCU activity. NLN OE increased _m_Ca^2+^ uptake under high Ca^2+^ conditions, coinciding with reduced MICU2 and increased MCU and MCUR1, a pattern opposite to *NLN* deficiency and consistent with MICU2-dependent control of channel gating. ATP23 OE enhanced uptake despite broadly reducing mtCU components, likely because MICU2, a major inhibitory gatekeeper, was markedly decreased. IMMP1L OE produced a Ca^2+^-dependent phenotype, increasing uptake at high Ca^2+^ but reducing uptake at low Ca^2+^, consistent with increased MICU1/MICU2-mediated gatekeeping near the activation threshold. In contrast, YME1L1 OE strongly suppressed _m_Ca^2+^ uptake and efflux, accompanied by reduced MCU, EMRE, MCUR1, and MICU3 and increased MICU1, MICU2, and MCUB. These coordinated changes predict impaired pore abundance, channel assembly, and gating. LACTB OE also suppressed _m_Ca^2+^ uptake and efflux despite MICU3 induction, likely reflecting reduced MCU abundance and broader impairment of transport capacity. Together, these observations indicate that IMS proteases regulate _m_Ca^2+^ homeostasis by selectively remodeling mtCU abundance and likely influencing complex organization rather than uniformly changing individual transport proteins. The CRC experiments indicate that IMS proteases contribute to mitochondrial resistance to Ca^2+^ overload. CRC integrates _m_Ca^2+^ uptake, efflux, matrix buffering, membrane potential maintenance, and resistance to mPTP opening. Thus, the widespread reduction in CRC across IMS protease KO models indicates that disruption of this protease network compromises multiple components of _m_Ca^2+^ homeostasis. Notably, CRC was reduced even in some models with increased low-Ca^2+^ uptake, suggesting that enhanced mtCU activity alone is insufficient to improve buffering capacity when NCLX function, matrix buffering, or mitochondrial integrity are not coordinately preserved. OE of nine of ten IMS proteases also reduced CRC, indicating that _m_Ca^2+^ buffering is sensitive to both protease loss and protease excess and that balanced IMS protease activity contributes to optimal resistance to Ca^2+^ overload.

In summary, this study defines IMS proteases as an integrated regulatory network that controls _m_Ca^2+^ transport by coordinating mtCU/NCLX abundance, protein stability, spatial organization, and function. Individual IMS proteases exert distinct but interconnected effects on _m_Ca^2+^ uptake, efflux, and buffering capacity. Future work should identify direct protease substrates, define how IMS proteases regulate mtCU assembly and turnover, and determine how these pathways are altered during physiological and pathological stress. Collectively, our findings provide a conceptual framework linking IMS proteostasis to _m_Ca^2+^ signaling and suggest that IMS protease pathways may represent potential therapeutic entry points in diseases characterized by disrupted _m_Ca^2+^ homeostasis.

## Methods

### Cell culture

HEK293T wild-type (WT) cells and IMS protease KO cell lines (*NLN^-/-^*, *ATP23^-/-^*, *IMMP1L^-/-^*, *IMMP2L^-/-^*, *YME1L1^-/-^*, *OMA1^-/-^*, *LACTB^-/-^*, *LACTB2^-/-^*, *PARL^-/-^*, and *HTRA2^-/-^*) were maintained in high-glucose Dulbecco’s Modified Eagle Medium (DMEM) supplemented with 10% FBS, 1 mM sodium pyruvate, 1% non-essential amino acids, and 1% penicillin–streptomycin. Details of all reagents and cell lines are provided in **Supplementary Table 4** and **Supplementary Table 5**, respectively. Cells were cultured at 37 °C in a humidified incubator containing 5% CO_2_.

### Immunoblotting

Cells were lysed in 1× RIPA lysis buffer supplemented with Halt™ Protease Inhibitor Cocktail. Cell suspensions were sonicated for 1 minute at 20% amplitude with 5-second pulses using a sonic dismembrator (Fisherbrand, Cat # FB120110). Lysates were centrifuged at 14,000 rpm for 20 minutes, and protein concentration was measured using the Pierce 660 nm Protein Assay. Equal amounts of protein (50 µg) were resolved by SDS–PAGE and transferred to PVDF membranes. Membranes were blocked in fluorescent blocking buffer for 1 hour at room temperature and incubated with primary antibodies overnight at 4 °C. Membranes were washed three times for 10 minutes each with TBS-T (TBS containing 0.1% Tween 20), incubated with the appropriate IRDye-conjugated secondary antibodies for 1 hour at room temperature, washed again, and visualized using a ChemiDoc imaging system. Signals were quantified using ImageJ. For OE studies, HEK293T cells were transiently transfected with empty pCMV vector or pCMV constructs encoding FLAG-tagged IMS proteases using FuGENE HD transfection reagent (Promega) according to the manufacturer’s protocol. Plasmids are listed in **Supplementary Table 6**. Twenty-four hours after transfection, cells were harvested and processed for western blot analysis. Antibody sources are provided in **Supplementary Table 7**. Details of all reagents are provided in **Supplementary Table 4.** Full-length, uncropped western blot images corresponding to the immunoblots presented in this study are provided in **Supplementary Figs. 6–11**. Densitometric fold-change values for mtCU proteins in IMS protease KO and OE conditions were analyzed in R. Protein and protease identifiers were standardized, values were converted to log_2_ fold changes, and changes ≥1.2-fold, ≤1/1.2-fold, or within this range were classified as increased, decreased, or unchanged, respectively. KO and OE profiles were visualized using spider radar plots, Sankey alluvial plots, bar plots, and dot plots.

### RNA isolation and global transcriptome sequencing

Total RNA was isolated from HEK293T control cells and IMS protease KO cell lines using the RNeasy Mini Kit (Qiagen, Cat #74104) according to the manufacturer’s instructions. RNA concentration and purity were determined spectrophotometrically using a Nanophotometer N50 (IMPLEN), and RNA integrity was assessed before library preparation. Only samples with an RNA integrity number (RIN) ≥8 were used for sequencing. Four independent biological replicates (n = 4) were analyzed for each genotype. RNA-seq libraries were prepared using the Illumina TruSeq Stranded mRNA Library Preparation Kit and sequenced on an Illumina NovaSeq 6000 platform to generate 150-bp paired-end reads, yielding an average depth of approximately 48 million reads per sample. Transcriptomic analyses were performed to identify gene-expression changes associated with loss of individual IMS proteases. Gene-expression data were analyzed in R Version 4.5.3. Zero or missing values were replaced with one-tenth of the minimum nonzero value within each sample column. Mean expression and log_2_ fold changes were calculated for each IMS protease KO group relative to HEK293T controls, and statistical significance was assessed using Welch’s *t*-test followed by Benjamini–Hochberg correction for multiple comparisons.

### Cycloheximide treatment and quantitative mitochondrial proteomics

To assess mitochondrial protein stability and turnover following acute inhibition of protein synthesis, HEK293T control cells and IMS protease KO cell lines were treated with cycloheximide (50 µg/mL) for 24 hours. Five independent biological replicates (n = 5) were analyzed for each genotype. After treatment, cells grown in 15-cm culture dishes were washed with ice-cold PBS and processed for mitochondrial isolation by differential centrifugation as described previously^72^. Briefly, cells were resuspended in ice-cold mitochondrial isolation buffer (MIB; 10 mM HEPES, pH 7.5, containing 200 mM mannitol, 70 mM sucrose, 1 mM EGTA, and protease inhibitor cocktail), homogenized using a Dounce homogenizer, and centrifuged at 500 × g for 10 minutes at 4 °C. Supernatants were collected and centrifuged at 12,000 × g for 15 minutes at 4 °C for crude mitochondrial pellets. Pellets were resuspended in MIB and washed twice by centrifuging at 12,000 × g for 20 minutes at 4 °C. Purified mitochondrial fractions were subjected to quantitative LC–MS/MS-based proteomic analysis to assess mitochondrial proteome changes caused by loss of individual IMS proteases under translational arrest. A total of 55 mitochondrial samples representing 11 experimental groups were processed, with each preparation containing approximately 400 µg protein in 50 µL. Samples were solubilized in SDS, heated, and quantified, after which 100 µg protein was supplemented with 14 pmol HSA internal standard and precipitated overnight with acetone. Protein pellets were reconstituted in Laemmli buffer, and 20 µg protein from each sample was electrophoresed approximately 1.5 cm into an SDS–PAGE gel. Individual lanes were excised, washed, reduced, alkylated, and digested overnight with 1 µg trypsin. Peptides were extracted using 50% acetonitrile, dried by SpeedVac, and reconstituted in 1% acetic acid. Samples were analyzed by data-independent acquisition on a Stellar mass spectrometer (ThermoFisher Scientific) using 5 m/z isolation windows across an m/z range of 350–950, with one full-scan spectrum acquired per cycle. DIA datasets were processed using DIA-NN and exported for downstream filtering and statistical analysis. Approximately 3,000 proteins were quantified, including at least 500 mitochondrial proteins.

Raw abundance data were processed in R Version 4.5.3, with zeros and non-detected values treated as missing. Proteins detected in fewer than two samples or absent from all experimental replicates were excluded. Missing values were imputed on the original scale using missForest, followed by log_2_ transformation^73^. IMS protease KO groups were compared with HEK293T controls using limma and TREAT (T-tests Relative to a Threshold) with a minimum 1.2-fold-change threshold, followed by Benjamini–Hochberg correction for multiple testing^74–76^. Positive and negative log_2_ fold changes were interpreted as relative stabilization and destabilization, respectively. Bubble plots, principal component analyses, Sankey alluvial plots, bar plots, and interaction networks were generated in R using protein log_2_ fold changes and associated statistical significance.

### Biotin-based proximity labeling and proteomics

HEK293T control cells and IMS protease KO cell lines were transfected with the corresponding control or IMS protease UltraID (UD) plasmids listed in **Supplementary Table 6**. For example, *OMA1^-/-^* cells were transfected with the OMA1-UD plasmid, whereas HEK293T control cells were transfected with the control UD plasmid. Four independent biological replicates (n = 4) were analyzed for each group. After transfection, cells were treated with 100 µM biotin for 30 minutes to induce proximity-dependent biotinylation. Cells were washed twice with ice-cold PBS, and mitochondria were isolated in MIB by differential centrifugation. Isolated mitochondria were lysed in UD lysis buffer containing 50 mM Tris, pH 7.4, 500 mM NaCl, 0.4% SDS, 2% Triton X-100, 1 mM dithiothreitol, and protease inhibitor cocktail. Cell suspensions were sonicated twice for 1 minute each at 20% amplitude with 5-second pulses using a sonic dismembrator. An equal volume of 50 mM Tris, pH 7.4, was then added, and lysates were centrifuged at 16,500 × g for 20 minutes. Supernatants were used for streptavidin-based immunoprecipitation (IP).

Streptavidin-enriched IP products were processed for LC–MS/MS-based quantitative proteomic analysis to identify IMS protease proximal interactomes^72^. Peptide mixtures were analyzed using an Orbitrap Eclipse mass spectrometer coupled to an Evosep Eno chromatography system (Evosep Biosystems, Odense, Denmark). For each sample, 500 ng peptides were loaded onto an Evotip, purified, and injected according to the manufacturer’s protocol. Peptides were separated on an EV1137 analytical column (15 cm × 150 µm, 1.5 µm) using a 45.4-minute linear gradient corresponding to the 30-samples-per-day method. Mobile phases consisted of water and acetonitrile, each containing 0.1% formic acid. Data-independent acquisition was performed using 44 static isolation windows, each 10 Th wide, across an *m/z* range of 400–900, with a cycle time of 3 seconds between adjacent survey spectra. Spectra were searched against the UniProt human protein FASTA database containing 20,395 annotated entries, downloaded in June 2021, using the Pulsar search engine in Spectronaut v20 (Biognosys AG, Switzerland). Search parameters included an FT-trap instrument setting, a precursor mass tolerance of 10 ppm, a monoisotopic fragment mass tolerance of 0.6 Da, full trypsin specificity, a maximum of two missed cleavages, and methionine oxidation (+15.995 Da) as a variable modification. This workflow enabled quantitative identification of proteins within the proximity-labeled interactomes of individual IMS proteases.

Raw protein-intensity data were processed in R. Proteins detected in fewer than two samples across the dataset were excluded, nonpositive intensities were treated as missing, and the remaining values were log_2_-transformed. Missing values were imputed using a left-censored minimum-probability method with a quantile threshold of 0.01, consistent with established approaches for low-abundance, missing-not-at-random proteomic measurements^57^. For each IMS protease condition, proteins with at least one experimentally observed value in the experimental group were retained and compared with the corresponding control using linear models and empirical Bayes variance moderation implemented in limma^55^. Robust empirical Bayes estimation with intensity-trend adjustment was applied, and log_2_ fold changes, nominal *P* values, and false-discovery-rate-adjusted *P* values were calculated. For proximity-interactome visualization, nonpositive log_2_ fold-change values were classified as not enriched or undetected, whereas positive enrichment values were retained for downstream bubble plot, Sankey alluvial plot, spider radar plot, and network analyses.

### Integrated analysis and plots

Integrated mtCU analysis was performed in R by harmonizing western blot KO/OE datasets, CHX stability proteomics, and UltraID proximity-proteomics datasets using common IMS protease and mtCU identifiers. Raw log_2_ fold changes were retained, whereas values were independently normalized for cross-dataset visualization. UltraID values ≤0 were treated as lacking positive enrichment, and missing measurements were displayed as hollow symbols in heatmaps and omitted from spider profiles. Radar plots, bubble plots, and bar plots were generated using ggplot2 in R.

### Evaluation of _m_Ca^2+^ flux

_m_Ca^2+^ flux was measured as previously described^28,46,72,77^. Briefly, _m_Ca^2+^ uptake and efflux were measured in permeabilized cells using Fura-FF^58^. For KO experiments, HEK293T control and IMS protease KO cells were used. For OE studies, HEK293T cells were transfected with either empty pCMV vector or pCMV vectors encoding FLAG-tagged IMS proteases (**Supplementary Table 6**) for 48 hours. Cells were harvested by trypsinization, counted using a TC20 Automated Cell Counter (Bio-Rad), and 2.5 × 10^6^ cells per condition were washed with intracellular medium (ICM; 120 mM KCl, 10 mM NaCl, 1 mM KH_2_PO_4_, and 20 mM HEPES-Tris, pH 7.2). Cells were then resuspended in ICM supplemented with succinate (2.5 mM), digitonin (25 µg), and thapsigargin (1.5 µM). Fura-FF (1 µM) was added immediately before analysis. Ca^2+^ flux was monitored using a PTI RatioMaster (Horiba). After baseline fluorescence was recorded for 300 seconds, CaCl_2_ (3 µM or 10 µM) was added to initiate _m_Ca^2+^ uptake. Ru360 (10 µM) and FCCP (10 µM) were subsequently added at 450 seconds and 600 seconds, respectively, to assess _m_Ca^2+^ uptake and efflux. Changes in extramitochondrial Ca^2+^ levels were continuously recorded throughout the experiment.

A custom MATLAB-based workflow (MathWorks, R2024a) was used to quantify _m_Ca^2+^ uptake and efflux kinetics from Fura-FF fluorescence traces. For uptake analysis, the maximal Fura-FF signal following CaCl_2_ addition at 300 seconds was identified, and the subsequent 50-second decay phase was fitted using a monoexponential function. For efflux analysis, the fluorescence response following Ru360 addition at 450 seconds was analyzed using an analogous monoexponential fit over the corresponding 50-second kinetic region. Uptake and efflux rates were calculated from the negative slopes of the log-transformed decay fits. For KO experiments, values were normalized to the mean of the corresponding HEK293T control replicates, whereas OE values were normalized to the mean of empty vector-transfected HEK293T control replicates. Fold changes were calculated relative to these respective baseline controls. MATLAB was also used to generate annotated traces for each replicate, displaying the complete Ca^2+^ flux profile and the kinetic regions used to calculate uptake and efflux rates. Control-normalized _m_Ca^2+^ uptake and efflux fold changes were analyzed in R across IMS protease KO and OE conditions at 3 µM and 10 µM Ca^2+^. IMS protease identifiers were standardized, and datasets were organized to compare KO versus OE profiles at each Ca^2+^ concentration and 3 µM versus 10 µM responses within each condition. Integrated spider plots were generated in Cartesian coordinates using ggplot2, with a fold change of 1 shown as the control reference.

### Measurement of _m_Ca^2+^ retention capacity

As published earlier^46,77^, CRC was measured in permeabilized cells using Fura-FF. HEK293T control and IMS protease KO cells, or HEK293T cells transfected with empty pCMV vector or FLAG-tagged IMS protease constructs (**Supplementary Table 6**) for 48 hours, were harvested, counted, and resuspended in ICM supplemented with succinate (2.5 mM), digitonin (25 µg), and thapsigargin (1.5 µM). Fura-FF was added immediately before analysis, and samples were measured using a PTI RatioMaster (Horiba). After baseline fluorescence was recorded for 300 seconds, 10 µM CaCl_2_ pulses were added every 50 seconds. Ca^2+^ additions continued until mitochondria could no longer sequester Ca^2+^, resulting in a sustained increase in extramitochondrial Ca^2+^ fluorescence. FCCP (10 µM) was then added to release accumulated _m_Ca^2+^ and confirm maximal Ca^2+^ loading. CRC was quantified by counting the number of Ca^2+^ boluses for which _m_Ca^2+^ uptake was ≥50% of the added calcium. The number of retained boluses for each condition was normalized to the corresponding control and expressed as a fold change.

### Statistics

All results are presented as mean ± SD. Statistical analyses were performed using GraphPad Prism 11.0.2 (GraphPad Software). All experiments were replicated at least three times, and measurements were obtained from distinct samples. Individual data points are shown in the figures with mean ± SD. Where appropriate, grouped analyses were performed using multiple unpaired *t*-tests or ordinary one-way ANOVA with multiple comparisons. For multiple unpaired *t*-tests, *P*-value thresholds were not corrected for multiple comparisons. For ordinary one-way ANOVA, the Gaussian distribution assumption was used, and the mean of each experimental column was compared with the mean of the corresponding control column. *P* values < 0.05 were considered statistically significant.

## Data and Resources Availability

The mass spectrometry proteomics datasets generated in this study have been deposited in the ProteomeXchange Consortium via the PRIDE partner repository. The cycloheximide-chase proteomics dataset is available under the project title “Mitochondria in HEK293 Gene KO Cells” with accession number PXD082181. The proximity-labeling proteomics dataset is available under the project title “Mapping IMS Proteases Interactome” with accession number PXD081788. All remaining data have been deposited in the figshare database and can be accessed using the DOI 10.6084/m9.figshare.33283668.

## Acknowledgements

We gratefully acknowledge Dr. Hadi Pourhadi and Dr. Jingyun Lee from the Proteomics and Metabolomics Shared Resource at Wake Forest University School of Medicine (WFUSM) for their assistance with the UltraID-based protein mass spectrometry experiments. We also thank Dr. Michael Kinter and Dr. Benjamin Miller from the Multiplexing Protein Analysis Core at the Oklahoma Medical Research Foundation (OMRF) for performing the proteomic mass spectrometry analyses associated with the cycloheximide-chase assays. We are grateful to all past and present members of the Tomar and Jadiya laboratories for their valuable scientific discussions, technical support, methodological assistance, and shared resources that contributed to this work. Language editing was assisted by Microsoft Copilot, and schematic illustrations were created using BioRender.com. Data analysis was performed using ImageJ, MATLAB 2024a, and R Version 4.5.3.

## Author Contributions

D.T. and P.J.: conceptualization, investigation, supervision, funding acquisition, and writing— review and editing. A.P.: data acquisition, data analysis, figure preparation, and writing—original draft. K.S.: formal analysis and visualization of transcriptomic and proteomic data and writing— methods. A.K.: data acquisition.

## Ethics Declarations

The authors declare no competing interests.

