## Supplementary for "Proteolytic control of mitochondrial calcium transport by intermembrane-space proteases"

**Running Title:** Proteolytic control of mitochondrial calcium flux

**Supplementary Figure 1. Validation and network-level analysis of IMS protease-dependent regulation of mtCU and NCLX proteins. A.** Representative western blots validating loss of individual IMS proteases in HEK293T cells, including *NLN*$^{-/-}$, *ATP23*$^{-/-}$, *IMMP1L*$^{-/-}$, *IMMP2L*$^{-/-}$, *YME1L1*$^{-/-}$, *OMA1*$^{-/-}$, *LACTB*$^{-/-}$, *LACTB2*$^{-/-}$, *PARL*$^{-/-}$, and *HTRA2*$^{-/-}$. **B.** Representative western blots confirming overexpression (OE) of Flag-tagged IMS proteases in HEK293 cells. **C.** Sankey alluvial plot generated from mtCU and NCLX protein-expression datasets in IMS protease KO and OE models. A concordant decrease indicates that both loss-of-function and OE reduce mtCU/NCLX protein abundance; a concordant increase indicates that both loss-of-function and OE increase mtCU/NCLX protein abundance. Loss-of-function-dominant regulation indicates a strong effect observed only under KO conditions, whereas OE-dominant regulation indicates a strong effect observed only under OE conditions. A reciprocal negative regulator indicates that loss of the IMS protease increases the protein while OE decreases it, whereas a reciprocal positive regulator indicates that loss of the IMS protease decreases the protein while OE increases it. **D.** Summary of the most strongly positively and negatively regulated mtCU/NCLX proteins in response to IMS protease KO and OE. **E.** Pearson correlation analysis showing coordinated changes in mtCU and NCLX protein abundance across IMS protease KO models. **F.** Pearson correlation analysis showing coordinated changes in mtCU/NCLX protein abundance across IMS protease OE models.

**Supplementary Figure 2.** IMS protease KO models display distinct mtCU/NCLX proteostasis signatures following cycloheximide (CHX) treatment. **A.** Principal component analysis (PCA) of CHX-based mitochondrial proteomics data showing the distribution of IMS protease KO models based on mtCU and NCLX protein-stability profiles. **B**. Directional analysis of mtCU and NCLX protein-stability changes across IMS protease KO models following CHX treatment. Stacked bars show the relative distribution of stabilized and destabilized mtCU/NCLX proteins in each KO condition. Stabilized proteins are shown in blue, whereas destabilized proteins are shown in red.

**Supplementary Figure 3. Integrated multidimensional analysis of IMS protease-dependent regulation of mtCU and NCLX proteins.** Dot plot summarizing quantitative values from four complementary datasets: mtCU/NCLX protein-expression changes in IMS protease KO models, IMS protease OE models, CHX-based mitochondrial proteomics, and UltraID proximity proteomics. This integrated visualization compares expression remodeling, protein-stability changes, and proximity enrichment across individual IMS proteases and mtCU/NCLX proteins.

**Supplementary Figure 4. Representative low-calcium $_{m}Ca^{2+}$ flux traces and uptake-to-efflux coupling in IMS protease KO cells. A–J.** Representative $_{m}Ca^{2+}$ flux traces from HEK293T control cells and IMS protease KO cell lines following acute stimulation with 3 μM $CaCl_2$. Traces show $_{m}Ca^{2+}$ uptake and subsequent efflux in *NLN*$^{-/-}$, *ATP23*$^{-/-}$, *IMMP1L*$^{-/-}$, *IMMP2L*$^{-/-}$, *YME1L1*$^{-/-}$, *OMA1*$^{-/-}$, *LACTB*$^{-/-}$, *LACTB2*$^{-/-}$, *PARL*$^{-/-}$, and *HTRA2*$^{-/-}$ cells compared with control cells. For representation, the same control trace is shown in panels A, B, D, E, G, H, I, and J. A second representative WT control trace is shown in panel C, and a third representative control trace is shown in panel F. **K.** Uptake-to-efflux relationship calculated from $_{m}Ca^{2+}$ flux assays performed with 10 μM $CaCl_2$. **L.** Uptake-to-efflux relationship calculated from $_{m}Ca^{2+}$ flux assays performed with 3 μM $CaCl_2$, summarizing the balance between $_{m}Ca^{2+}$ entry and extrusion under low-calcium conditions. **M.** Integrated radar plot comparing normalized $_{m}Ca^{2+}$ uptake and efflux rates across all IMS protease KO models under 10 μM and 3 μM $CaCl_2$ stimulation. Blue traces represent the 10 μM $CaCl_2$ condition, and red traces represent the 3 μM $CaCl_2$ condition, illustrating the distinct $Ca^{2+}$ transport signatures associated with each IMS protease deficiency across high and low $Ca^{2+}$ loads. All quantitative analyses were performed using all independent control and KO biological replicate traces, including the representative traces shown in panels A–J, and were not restricted to the representative traces displayed in the figure. All flux quantifications were performed from at least three independent biological replicates. Data are presented as mean ± SD. Statistical significance was determined using ordinary one-way ANOVA with multiple comparisons; $^{*}p < 0.05$, $^{**}p < 0.01$, $^{****}p < 0.0001$, ns = not significant ($p \geq 0.05$).

**Supplementary Figure 5. Representative low-calcium $_{m}Ca^{2+}$ flux traces and uptake-to-efflux coupling following IMS protease OE. A–J.** Representative $_{m}Ca^{2+}$ flux traces from HEK293T control cells and cells overexpressing individual IMS proteases following acute stimulation with 3 μM $CaCl_2$. Traces show $_{m}Ca^{2+}$ uptake and subsequent efflux in cells overexpressing NLN, ATP23, IMMP1L, IMMP2L, YME1L1, OMA1, LACTB, LACTB2, PARL, or HTRA2 compared with control cells. For representation in panels A–J, the same control trace is shown alongside each IMS protease OE trace. **K.** Uptake-to-efflux relationship calculated from $_{m}Ca^{2+}$ flux assays performed with 10 μM $CaCl_2$. **L.** Uptake-to-efflux relationship calculated from $_{m}Ca^{2+}$ flux assays performed with 3 μM $CaCl_2$, summarizing the balance between $_{m}Ca^{2+}$ entry and extrusion under low-$Ca^{2+}$ conditions following IMS protease OE. **M.** Integrated radar plot comparing normalized $_{m}Ca^{2+}$ uptake and efflux rates across all IMS protease OE models under 10 μM and 3 μM $CaCl_2$ stimulation. Blue traces represent the 10 μM $CaCl_2$ condition, and red traces represent the 3 μM $CaCl_2$ condition, illustrating the distinct $Ca^{2+}$ transport signatures associated with overexpression of individual IMS proteases across high and low $Ca^{2+}$ loads. All quantitative analyses were performed using all independent control and IMS protease OE biological replicate traces, including the representative traces shown in panels A–J, and were not restricted to the representative traces displayed in the figure. All flux quantifications were performed from at least three independent biological replicates. Data are presented as mean ± SD. Statistical significance was determined using one-way ANOVA with multiple comparisons; ***$p < 0.001$, ****$p < 0.0001$, ns = not significant ($p \geq 0.05$).

**Supplementary Figures 6–11. Full-length western blots supporting IMS protease and mtCU/NCLX protein analyses.** Full-length western blot images are shown for the corresponding cropped blot panels presented in the main figures.

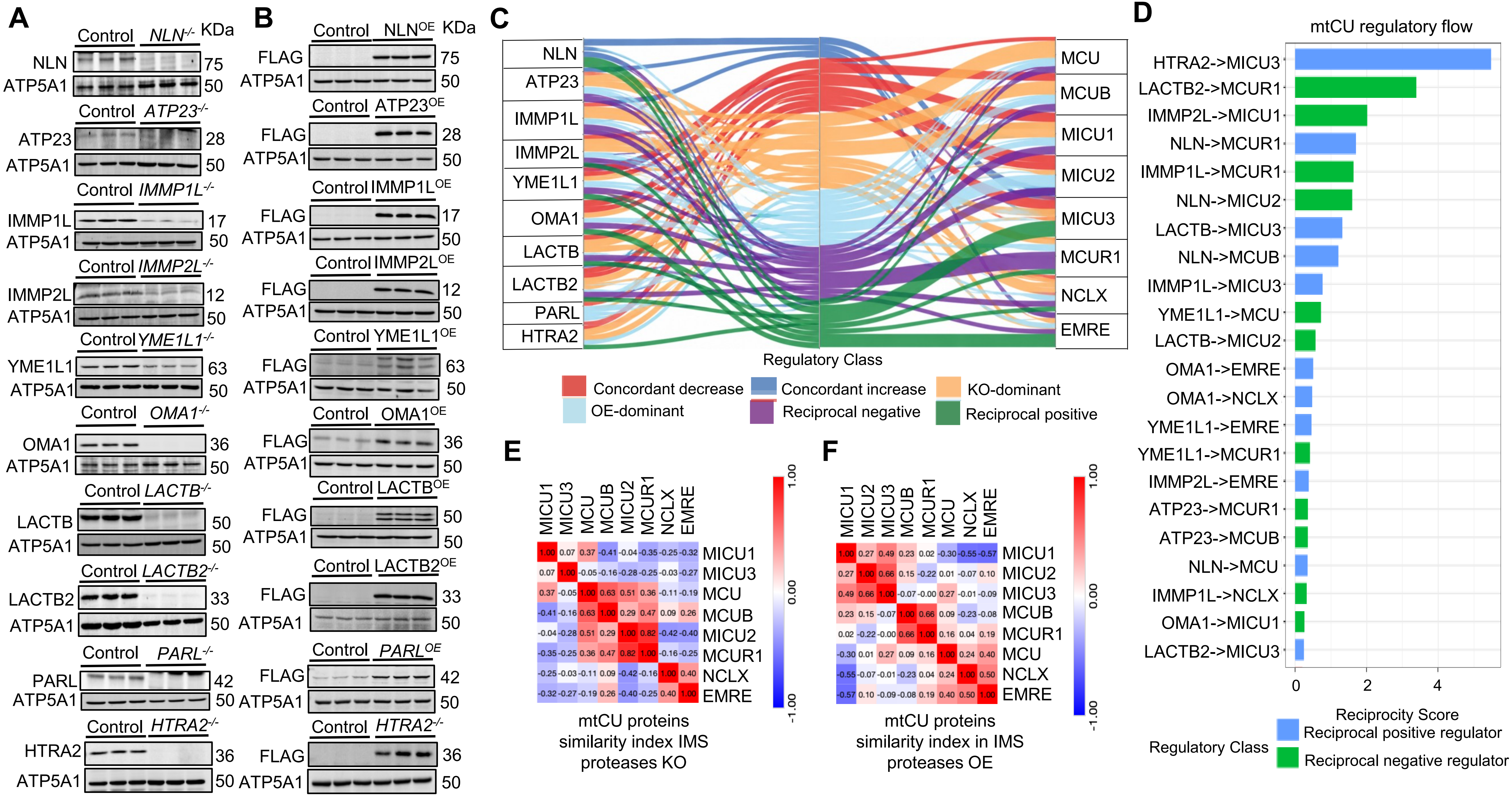


**Supplementary Figure 1**

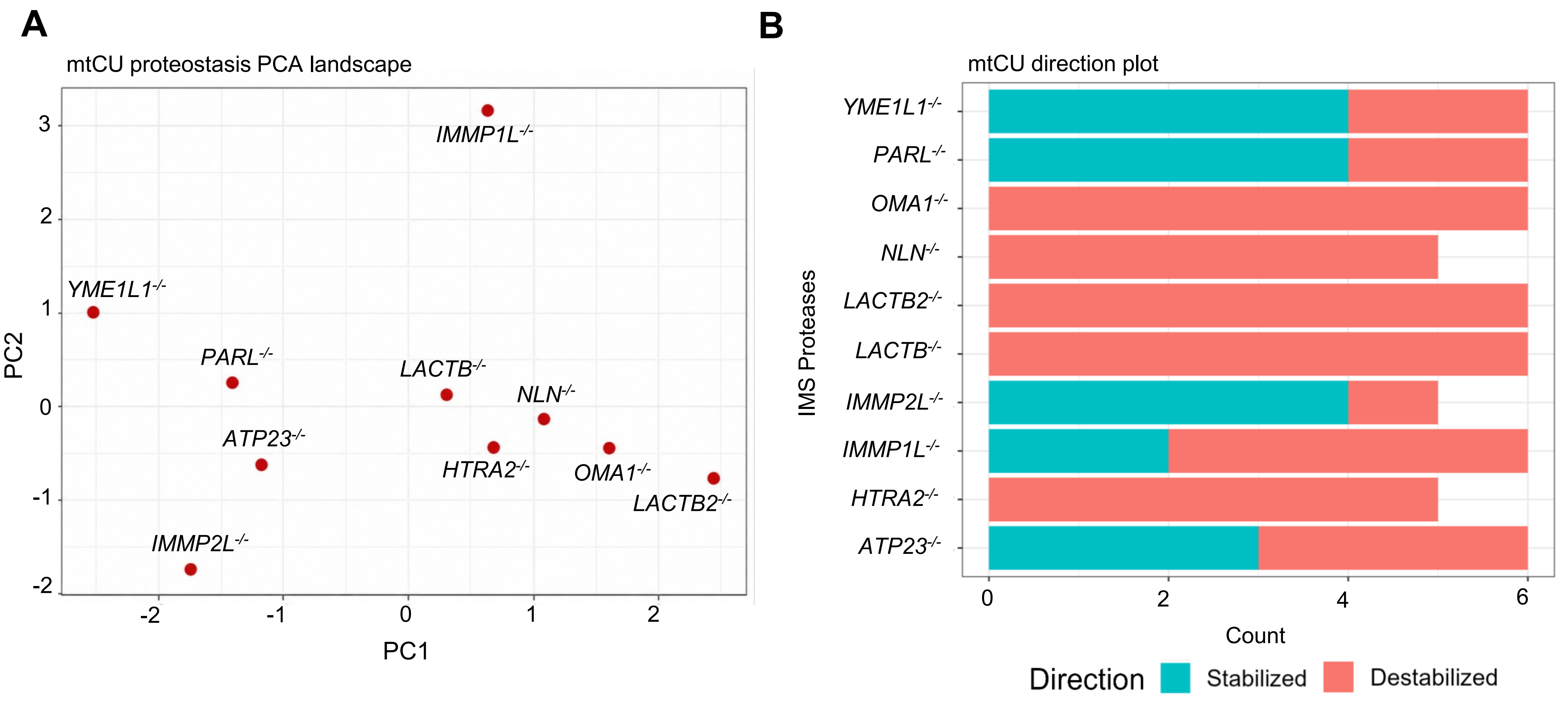


Each IMS protease KO generated a distinct mtCU proteostasis landscape

Individual proteases exerted unique stabilizing and destabilizing effects on specific mtCU proteins

**Supplementary Figure 2**

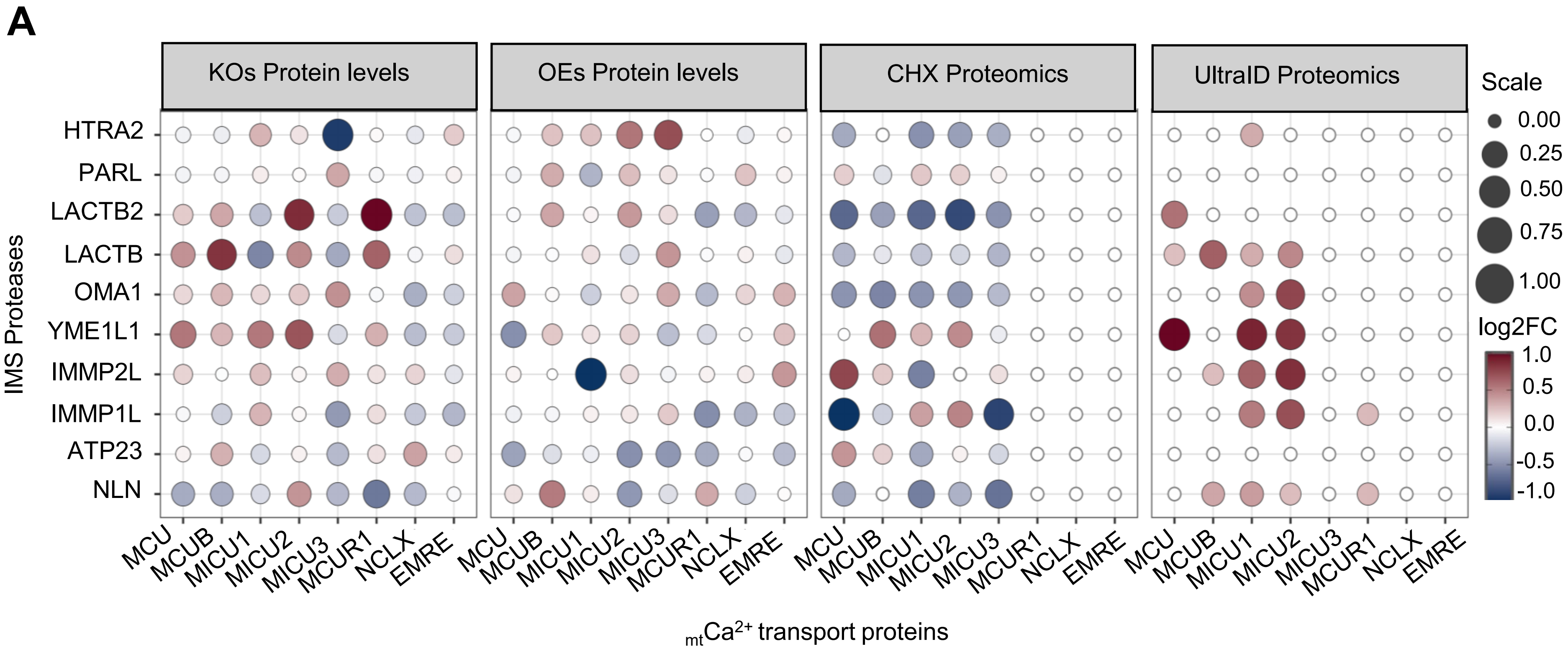


**Supplementary Figure 3**

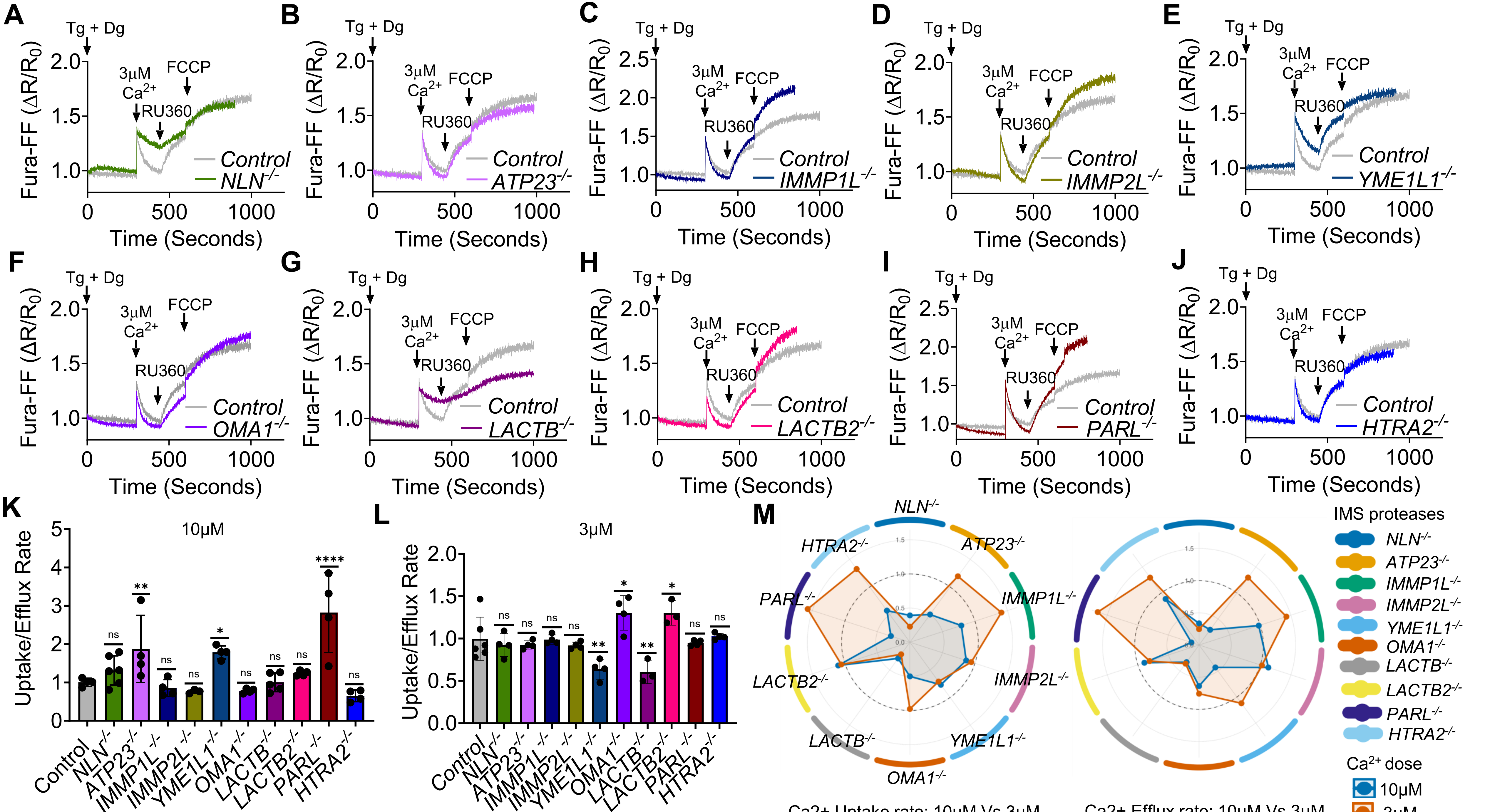


**Supplementary Figure 4**

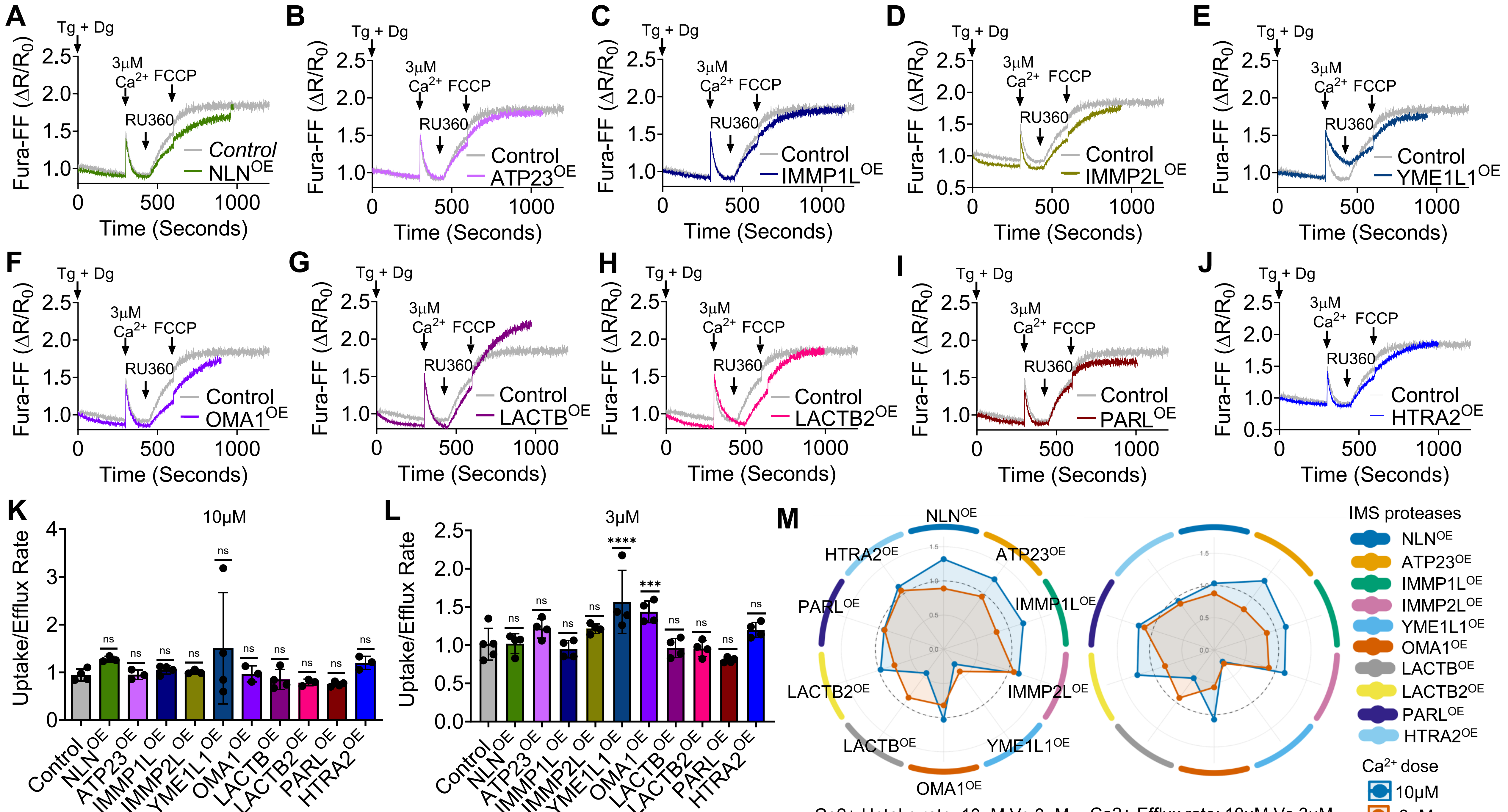


**Supplementary Figure 5**

Full-length blots for Figure 1A

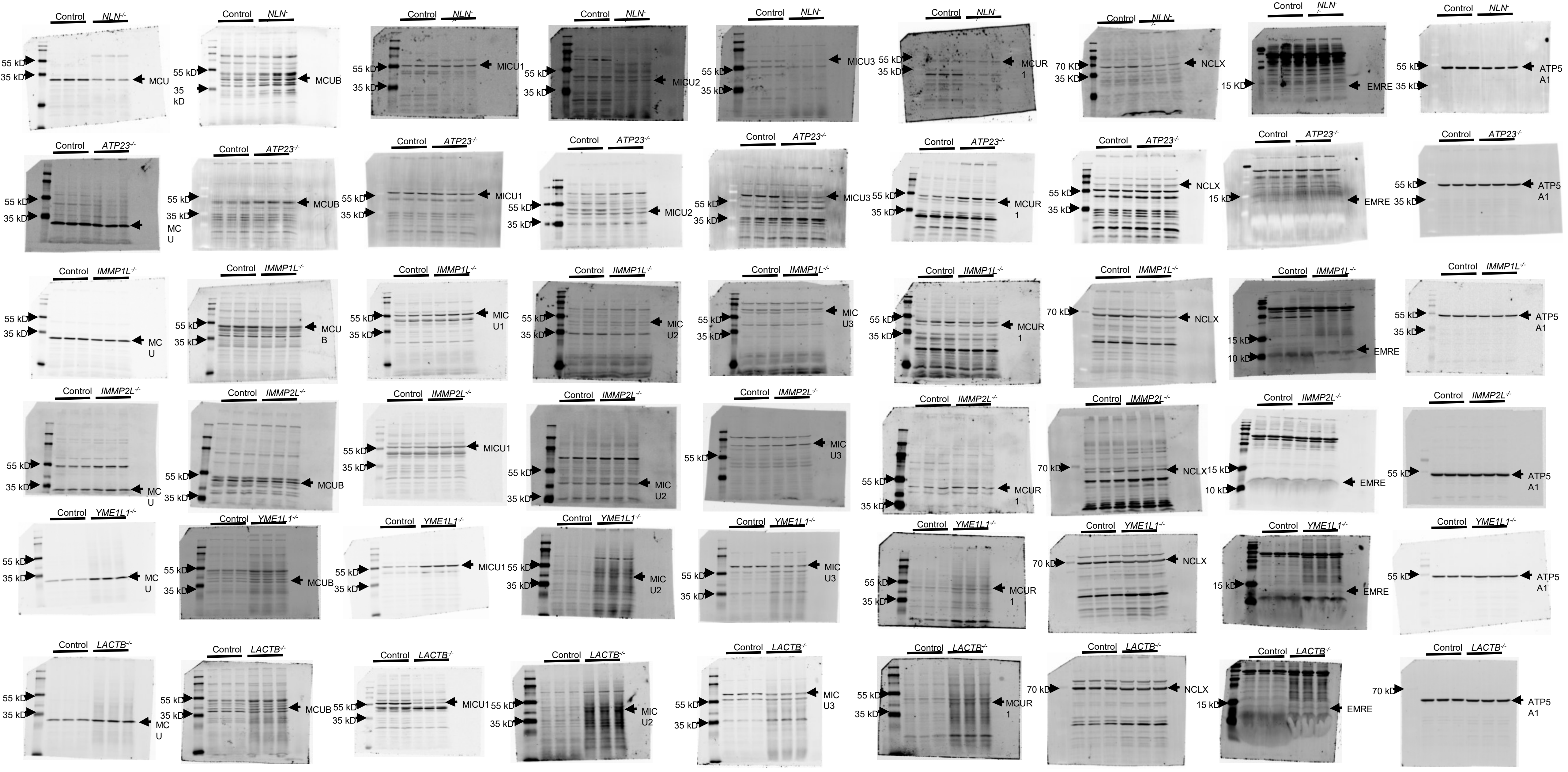


#### Supplementary Figure 6

Full-length blots for Figure 1A Continued

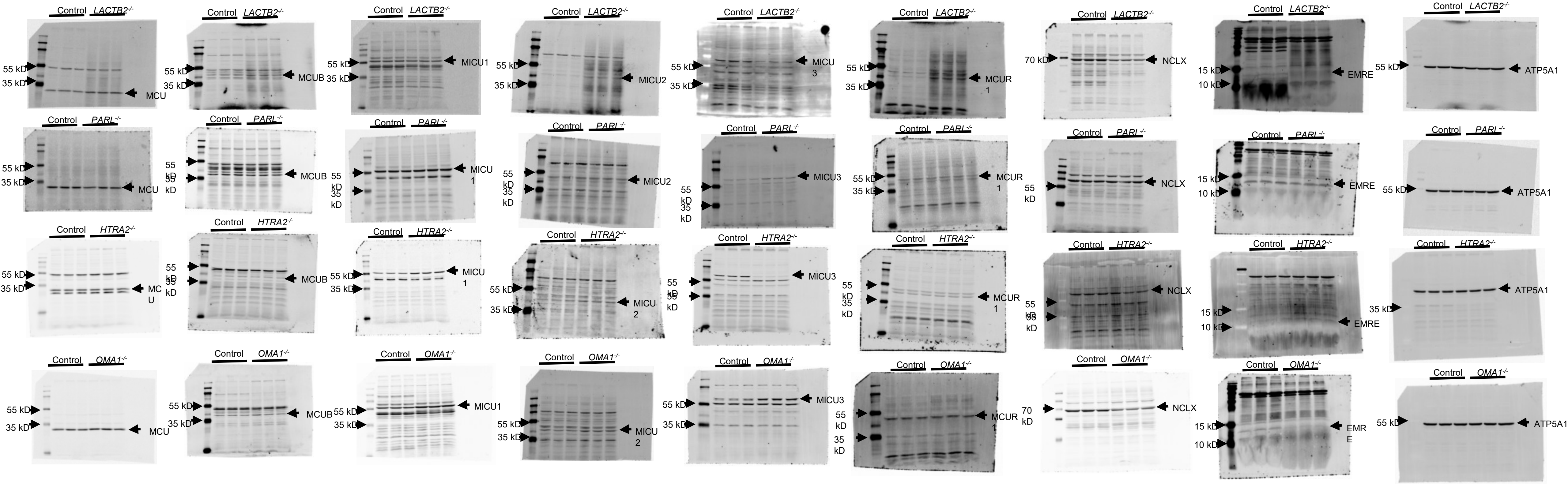


### Supplementary Figure 7

Full-length blots for Figure 1B

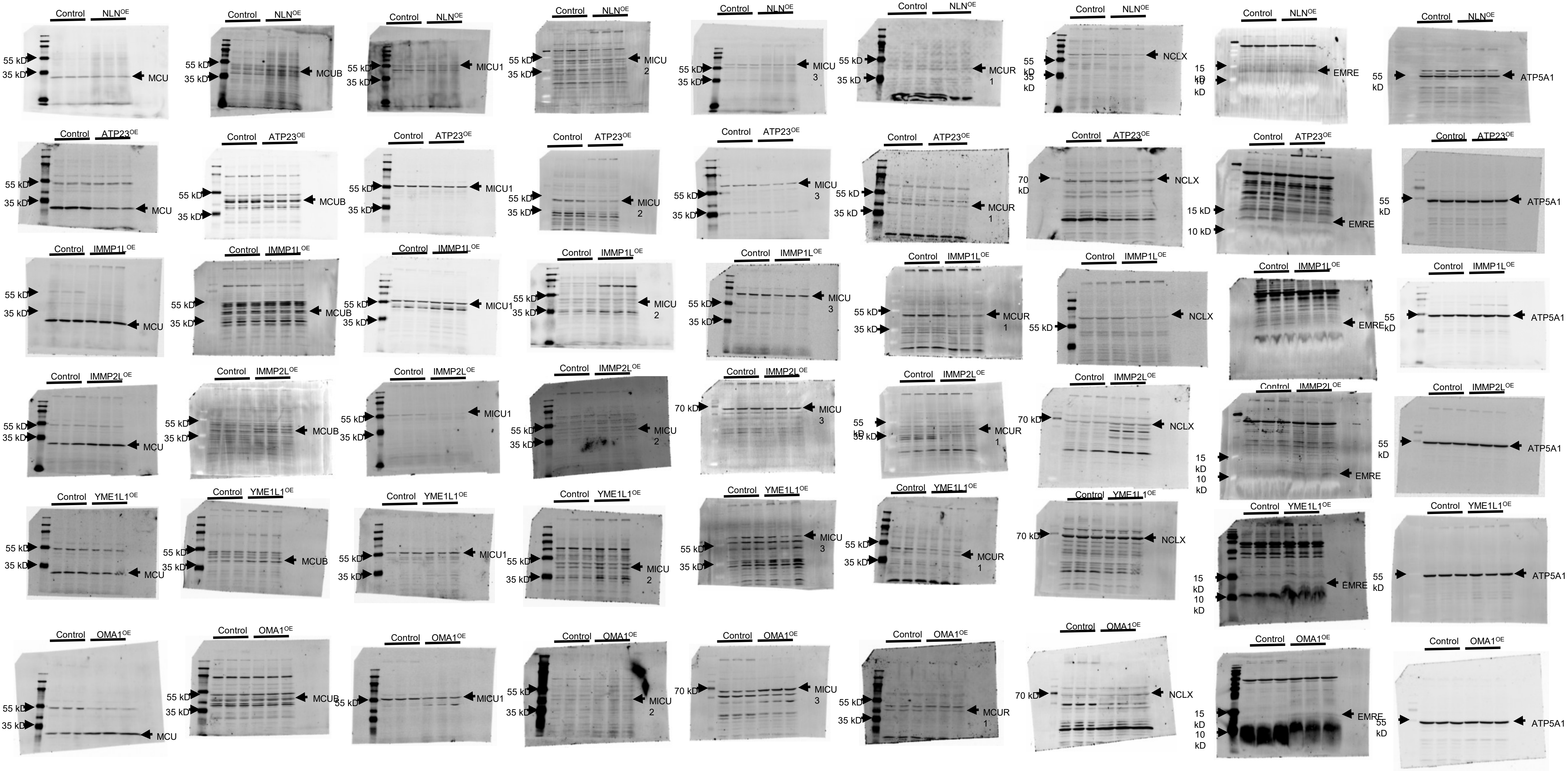


### Supplementary Figure 8

Full-length blots for Figure 1B Continued

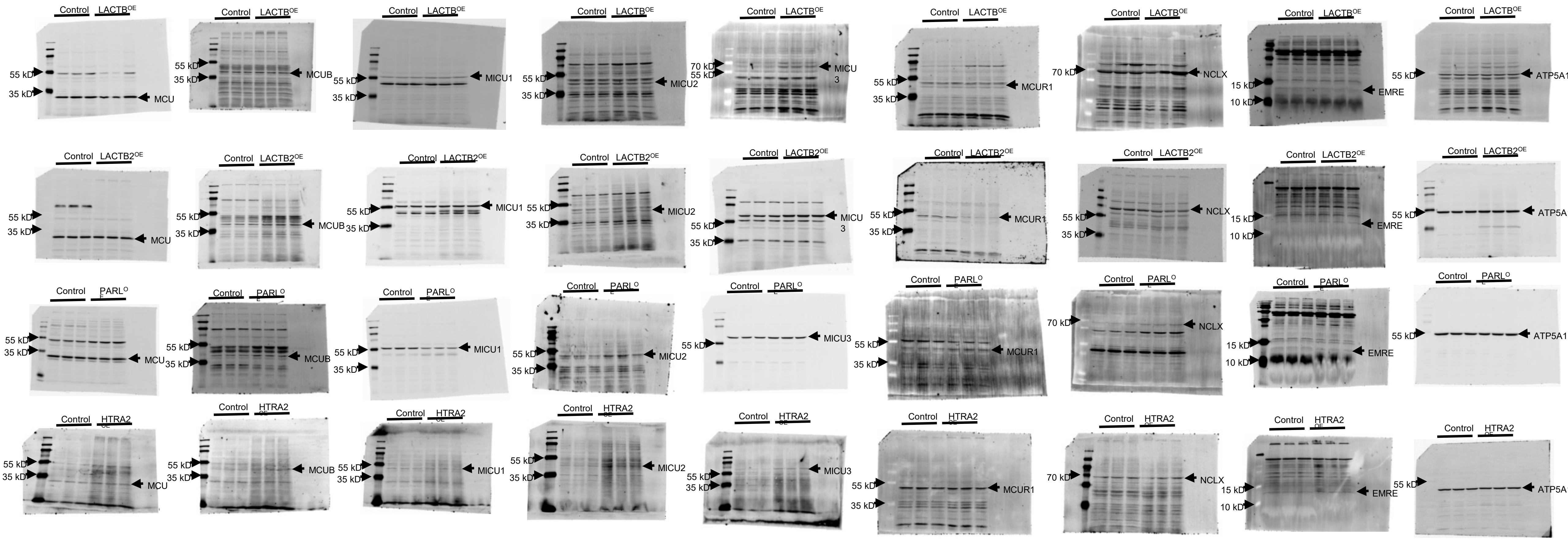


### Supplementary Figure 9

Full-length blots for Supplementary Figure 1A

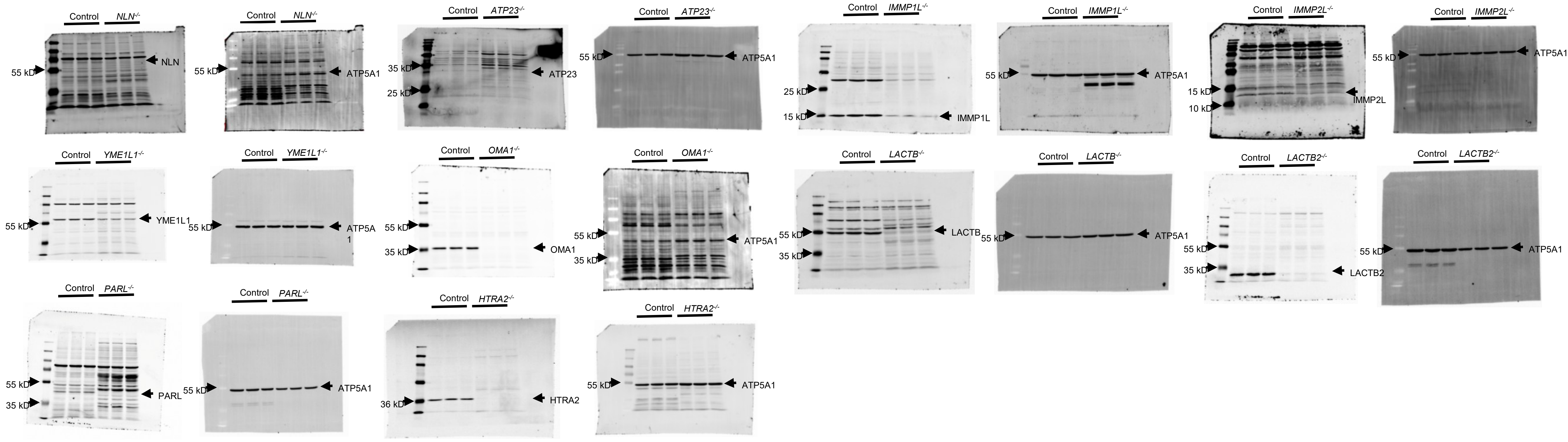


### Supplementary Figure 10

Full-length blots for Supplementary Figure 1B

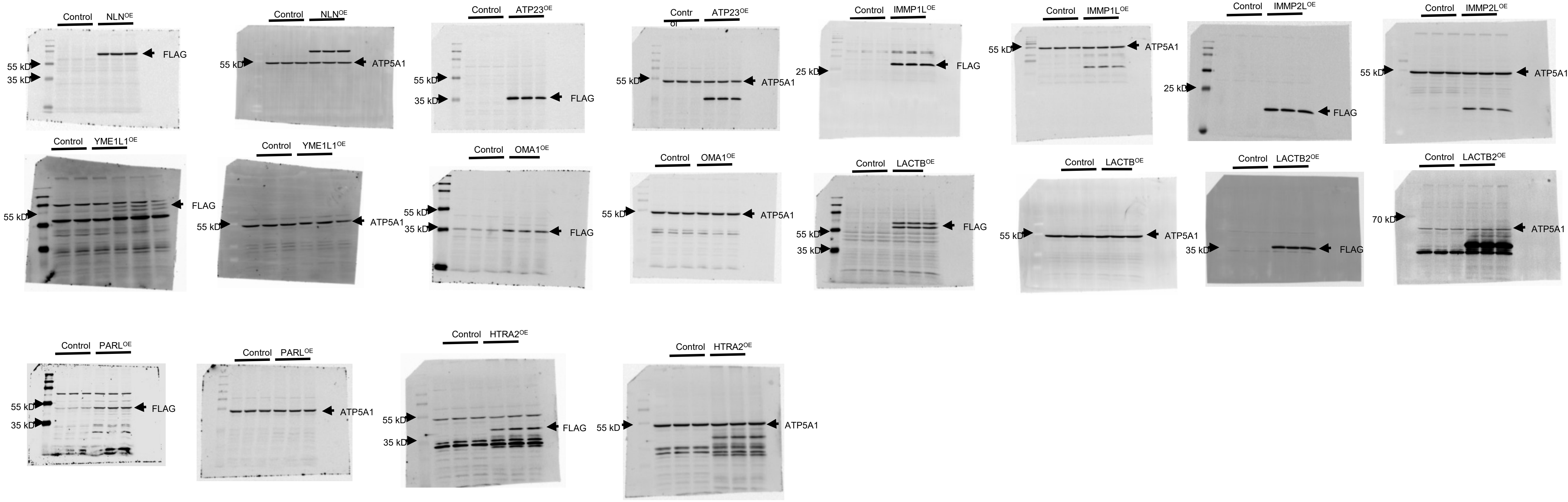


### Supplementary Figure 11

#### Supplementary Tables

| mtCU | IMS Proteases | | | | | | | | | |
|---|---|---|---|---|---|---|---|---|---|---|
| | *NLN*$^{-/-}$ | *ATP23*$^{-/-}$ | *IMMP1L*$^{-/-}$ | *IMMP2L*$^{-/-}$ | *YME1L1*$^{-/-}$ | *OMA1*$^{-/-}$ | *LACTB*$^{-/-}$ | *LACTB2*$^{-/-}$ | *PARL*$^{-/-}$ | *HTRA2*$^{-/-}$ |
| MCU | 0.003538 | 0.108231 | 0.508645 | 0.002252 | 0.000004 | 0.001877 | 0.000746 | 0.011576 | 0.31389 | 0.298 |
| MCUB | 0.057474 | 0.004416 | 0.002394 | 0.651648 | 0.00106 | 0.011906 | 0.001182 | 0.002615 | 0.28331 | 0.074 |
| MICU1 | 0.01648 | 0.000163 | 0.005939 | 0.002405 | 0.000055 | 0.000092 | 0.001419 | 0.004526 | 7.5E-05 | 0.0001 |
| MICU2 | 0.000588 | 0.041769 | 0.693324 | 0.271226 | 0.000227 | 0.005593 | 0.000071 | 0.000206 | 0.77082 | 0.021 |
| MICU3 | 0.001924 | 0.000249 | 0.059347 | 0.014362 | 0.033238 | 0.002103 | 0.000399 | 0.009146 | 0.00065 | 0.0001 |
| MCUR1 | 0.000105 | 0.131447 | 0.036628 | 0.341245 | 0.001505 | 0.070887 | 0.000007 | 0.000119 | 0.1528 | 0.038 |
| NCLX | 0.006167 | 0.000211 | 0.004418 | 0.02466 | 0.003664 | 0.000098 | 0.431901 | 0.002618 | 0.03719 | 0.061 |
| EMRE | 0.605361 | 0.157814 | 0.004309 | 0.055949 | 0.011536 | 0.000148 | 0.004059 | 0.000031 | 0.01899 | 0.001 |

Significantly Upregulated

Significantly Downregulated

**Supplementary Table 1. Statistical significance of mtCU/NCLX protein-expression changes in IMS protease KO cells.** Protein abundance was quantified from western blots using ImageJ, normalized to the corresponding loading control, and compared with HEK293T control cells. Statistical significance was determined using multiple unpaired t-tests from three independent replicates. Statistically significant changes were defined as $p < 0.05$ and are highlighted in the table. Blue highlights indicate significant downregulation, whereas red highlights indicate significant upregulation.

| mtCU | IMS Proteases | | | | | | | | | |
|---|---|---|---|---|---|---|---|---|---|---|
| | NLN$^{OE}$ | ATP23$^{OE}$ | IMMP1L$^{OE}$ | IMMP2L$^{OE}$ | YME1L1$^{OE}$ | OMA1$^{OE}$ | LACTB$^{OE}$ | LACTB2$^{OE}$ | PARL$^{OE}$ | HTRA2$^{OE}$ |
| MCU | 0.081886 | 0.008679 | 0.334469 | 0.253498 | 0.000271 | 0.000328 | 0.202907 | 0.671389 | 0.161952 | 0.495713 |
| MCUB | 0.000442 | 0.010508 | 0.617615 | 0.166388 | 0.026092 | 0.421843 | 0.713066 | 0.000569 | 0.000681 | 0.005944 |
| MICU1 | 0.027989 | 0.086607 | 0.504991 | 0.000017 | 0.31215 | 0.004685 | 0.000773 | 0.441196 | 0.002413 | 0.000396 |
| MICU2 | 0.00402 | 0.004437 | 0.012663 | 0.041776 | 0.012875 | 0.071788 | 0.004637 | 0.001668 | 0.000725 | 0.001568 |
| MICU3 | 0.001857 | 0.002519 | 0.256086 | 0.096603 | 0.052732 | 0.000384 | 0.003267 | 0.007192 | 0.00104 | 0.010292 |
| MCUR1 | 0.005012 | 0.002889 | 0.000751 | 0.050046 | 0.186247 | 0.015142 | 0.201753 | 0.002726 | 0.877253 | 0.927948 |
| NCLX | 0.031731 | 0.711795 | 0.005984 | 0.027429 | 0.555226 | 0.129653 | 0.244418 | 0.0004 | 0.005858 | 0.339526 |
| EMRE | 0.87127 | 0.000763 | 0.020339 | 0.006424 | 0.000323 | 0.001109 | 0.008988 | 0.184362 | 0.35636 | 0.751995 |

Significantly Upregulated

Significantly Downregulated

**Supplementary Table 2. Statistical significance of mtCU and NCLX protein-expression changes in IMS protease OE cells.** Protein abundance was quantified from western blots using ImageJ, normalized to the corresponding loading control, and compared with HEK293T control cells. Statistical significance was determined using multiple unpaired t-tests from three independent replicates. Statistically significant changes were defined as $p < 0.05$ and are highlighted in the table. Blue highlights indicate significant downregulation, whereas red highlights indicate significant upregulation.

| mtCU | IMS Proteases | | | | | | | | | |
|---|---|---|---|---|---|---|---|---|---|---|
| | *NLN*$^{-/-}$ | *ATP23*$^{-/-}$ | *IMMP1L*$^{-/-}$ | *IMMP2L*$^{-/-}$ | *YME1L1*$^{-/-}$ | *OMA1*$^{-/-}$ | *LACTB*$^{-/-}$ | *LACTB2*$^{-/-}$ | *PARL*$^{-/-}$ | *HTRA2*$^{-/-}$ |
| *MCU* | 0.899028 | 0.211079 | 0.82114554 | 0.01479594 | 0.09714818 | 0.026382 | 0.228429 | 0.042099 | 0.220767 | 0.019722 |
| *MCUB* | 0.856195 | 0.785891 | 0.60843782 | 0.20095688 | 0.16029287 | 0.200734 | 0.607006 | 0.489436 | 0.67015 | 0.086451 |
| *MICU1* | 0.926574 | 0.386063 | 0.46487803 | 0.0203272 | 0.22481511 | 0.005503 | 0.394061 | 0.097469 | 0.477497 | 0.061192 |
| *MICU2* | 0.921787 | 0.460873 | 0.7514559 | 0.04351945 | 0.53886599 | 0.197302 | 0.256347 | 0.064153 | 0.057788 | 0.023853 |
| *MICU3* | 0.831851 | 0.908176 | 0.46761856 | 0.45274984 | 0.73911353 | 0.892819 | 0.979874 | 0.437935 | 0.588925 | 0.34774 |
| *MCUR1* | 0.831851 | 0.975714 | 0.16653681 | 0.03311889 | 0.03154487 | 0.034526 | 0.559993 | 0.011626 | 0.175363 | 0.03771 |
| *NCLX* | 0.9979 | 0.23898 | 0.9445197 | 0.7193065 | 0.60617894 | 0.768412 | 0.503877 | 0.185632 | 0.750812 | 0.048482 |
| *EMRE* | 0.831851 | 0.863308 | 0.7713751 | 0.67223412 | 0.09458035 | 0.246003 | 0.791301 | 0.446421 | 0.068578 | 0.00209 |

Significantly Upregulated

Significantly Downregulated

**Supplementary Table 3. Adjusted *p*-values for differential expression of mtCU complex components and NCLX in IMS protease KO cells.** This table provides the adjusted *p*-values corresponding to the differential gene-expression data shown in the heatmap in Figure 2A. Statistically significant changes were defined as adjusted *p*-values (*padj*) < 0.05 and are highlighted in the table. Blue shading indicates significantly decreased gene expression, whereas red shading indicates significantly increased gene expression relative to wild-type cells. Non-highlighted cells denote genes whose expression was not significantly altered.

**Supplementary Table 4**

| Chemicals | Source | Identifier (Catalogue #) |
|---|---|---|
| Digitonin | Sigma-Aldrich | 300410-1GM |
| Sodium Succinate Dibasic Hexahydrate | Sigma-Aldrich | S2378-100G |
| Thapsigargin | Millipore Sigma | 586005-1MG |
| FuraFF | Cayman Chemicals | 20415 |
| FCCP | Cayman Chemicals | 15218 |
| Calcium chloride | Thermo Fisher Scientific | J63122.AE |
| Ru360 | Sigma-Aldrich | 557440-500UG |
| 1X DPBS | Thermo Fisher Scientific | 14190144 |
| 10X RIPA lysis buffer | Millipore Sigma | 20-188 |
| Halt™ Phosphatase Inhibitor Cocktail | Thermo Fisher Scientific | 78420 |
| Halt™ Protease Inhibitor Cocktail | Thermo Fisher Scientific | 78429 |
| Fluorescent Blocking Buffer | Rockland | MB-070 |
| SDS, 20% Solution | Research Products International (RPI) | L23100-500.0 |
| TEMED | Fisher Scientific | BP150-20 |
| Ammonium persulfate | Fisher Bioreagents | BP179100 |
| Tween 20 | Sigma Aldrich | P9416 |
| Protogel | National Diagnostics | EC-890 |
| 4X ProtoGel Resolving Buffer | National Diagnostics | EC-892 |
| Methanol | Spectrum Chemical | M12404LTGLCS4 |
| Tris Base | Fisher Bioreagents | BP-152-1 |
| Pierce 660nm Protein assay reagent | Thermo Fisher Scientific | 22660 |
| Tris-Glycine-SDS, 10X Solution | Fisher Scientific | BP1341-1 |
| Tris-Glycine, 10X Solution | Fisher Scientific | BP1306-1 |
| Laemmli SDS sample buffer, reducing (6X) | Thermo Fisher Scientific | J61337.AD |
| PageRuler™ Plus Prestained Protein Ladder | Fisher Scientific | PI26619 |
| Immobilon transfer membrane | Millipore Sigma | IPFL00010 |
| D-Mannitol | ACROS Organics | 125345000 |
| Sucrose | MP Biomedicals | 821713 |
| EGTA | Millipore Sigma | 324626 |
| Triton-X 100 | Promega | HF-141 |
| Chelex 100 Chelating Resin | Bio-Rad | 422822 |
| Potassium Chloride | Fisher Scientific | BP366-500 |
| Sodium chloride | Millipore Sigma | S9888 |
| Potassium phosphate monobasic | Millipore Sigma | P5655-500G |
| HEPES Buffer | Corning | 25-060-CI |
| Biotin | Sigma-Aldrich | B4501-1G |

| | | |
|---|---|---|
| Dynabeads™ MyOne™ Streptavidin T1 | Thermo Fisher Scientific | 65602 |
| Dithiothreitol (DTT) | Thermo Fisher Scientific | 18091050 |
| HyClone™ Dulbecco's Modified Eagle Medium (DMEM) with high glucose | Cytiva | SH30022.02 |
| Opti-MEM | Gibco | 31985-070 |
| Sodium Pyruvate | Corning | 25-000-CI |
| MEM Nonessential Amino Acids | Corning | 25-025-CI |
| Penicillin-Streptomycin Solution | Corning | 30-002-CI |
| FuGENE® HD Transfection Reagent | Promega | E2311 |
| Fetal Bovine Serum (FBS) | Alkali Scientific | 502335958 |
| RNeasy Mini Kit | Qiagen | 74104 |

**Supplementary Table 4. Reagents used in this study.**
This table provides a comprehensive list of reagents used throughout the study, including chemicals, biological reagents, assay kits, and other experimental materials. The source/vendor and corresponding catalog number are listed for each reagent.

**Supplementary Table 5**

| Cell lines | Source | Identifier (Catalogue #) |
|---|---|---|
| HEK293T WT | Ubigene | YC-A006 |
| HEK293T *ATP23*$^{-/-}$ | Ubigene | YKO-HT24701 |
| HEK293T *IMMP1L*$^{-/-}$ | Ubigene | YKO-HT26539 |
| HEK293T *IMMP2L*$^{-/-}$ | Ubigene | YKO-HT24066 |
| HEK293T *OMA1*$^{-/-}$ | Ubigene | YKO-HT25030 |
| HEK293T *PARL*$^{-/-}$ | Ubigene | CK24-041-A4 (Custom made) |
| HEK293T *HTRA2*$^{-/-}$ | Ubigene | CK24-042-A5 (Custom made) |
| HEK293T *LACTB2*$^{-/-}$ | Ubigene | YKO-HT21041 |
| HEK293T *NLN*$^{-/-}$ | Ubigene | YKO-HT22560 |
| HEK293T *YME1L1*$^{-/-}$ | Ubigene | YKO-HT19262 |
| HEK293T *LACTB*$^{-/-}$ | Ubigene | YKO-HT24958 |

**Supplementary Table 5. Cell lines used in this study.** This table lists all cell lines used in the study, including wild-type HEK293T cells and individual IMS protease KO lines, together with their source and catalog or identifier information.

**Supplementary Table 6**

| Plasmids | Source | Identifier (Catalogue #) |
|---|---|---|
| PCMV-NLN-FLAG-MYC | OriGene | RC212447 |
| PCMV-ATP23-FLAG-MYC | OriGene | RC203623 |
| PCMV-IMMP1L-FLAG-MYC | OriGene | RC204909 |
| PCMV-IMMP2L-FLAG-MYC | OriGene | RC215438 |
| PCMV-YME1L1-FLAG-MYC | OriGene | RC203167 |
| PCMV-OMA1-FLAG-MYC | OriGene | RC204507 |
| PCMV-LACTB-FLAG-MYC | OriGene | RC209605 |
| PCMV-LACTB2-FLAG-MYC | OriGene | RC201465 |
| PCMV-PARL-FLAG-MYC | OriGene | RC204096 |
| PCMV-HTRA2-FLAG-MYC | OriGene | RC222206 |
| pCMV6-AC-HA | OriGene | PS100004 |
| pRP[Exp]-Bsd-CMV NLN-Ultra ID | Vector Builder | VB250410-1393xab |
| pRP[Exp]-Bsd-CMV ATP23-Ultra ID | Vector Builder | VB250410-1400mwp |
| pRP[Exp]-Bsd-CMV IMMP1L-Ultra ID | Vector Builder | VB250410-1396npu |
| pRP[Exp]-Bsd-CMV IMMP2L-Ultra ID | Vector Builder | VB250410-1389wsr |
| pRP[Exp]-Bsd-CMV YME1L1-Ultra ID | Vector Builder | VB250410-1401bdc |
| pRP[Exp]-Bsd-CMV LACTB-Ultra ID | Vector Builder | VB250410-1395neu |
| pRP[Exp]-Bsd-CMV LACTB2-Ultra ID | Vector Builder | VB250410-1402vvx |
| pRP[Exp]-Bsd-CMV PARL-Ultra ID | Vector Builder | VB250410-1399hpa |
| pRP[Exp]-Bsd-CMV OMA1-Ultra ID | Vector Builder | VB250410-1397sys |
| pRP[Exp]-Bsd-CMV HTRA2-Ultra ID | Vector Builder | VB250410-1391bbr |
| pRP[Exp]-Bsd-CMV Control Ultra ID | Vector Builder | VB250421-1197cam |

**Supplementary Table 6. Plasmids used in this study.** This table provides details for all plasmid constructs used in the study, including the vector backbone, encoded gene or protein, epitope tag, source, and catalog or identifier information.

133 **Supplementary Table 7**

| Antibodies | Dilution | Source | Identifier (Catalogue #) |
|---|---|---|---|
| Anti NLN Rabbit pAb | 1:200 | Proteintech | 14763-1-AP |
| Anti-XRCC6BP1 (ATP23) Rabbit pAb | 1:500 | Proteintech | 16076-1-AP |
| Anti IMMP1L Rabbit pAb | 1:500 | Proteintech | PA524322 |
| Anti IMMP2L Rabbit pAb | 1:500 | Proteintech | 15970-1-AP |
| Anti YME1L1 Rabbit pAb | 1:1000 | Proteintech | 11510-1-AP |
| Anti OMA1 Rabbit pAb | 1:1000 | Proteintech | 17116-1-AP |
| Anti LACTB Mouse pAb | 1:1000 | Proteintech | 66785-1-Ig |
| Anti LACTB2 Rabbit pAb | 1:1000 | Proteintech | 16783-1-AP |
| Anti PARL Rabbit pAb | 1:500 | Proteintech | 26679-1-AP |
| Anti Htra2/Omi Rabbit pAb | 1:1000 | Cell Signaling Technology | 9745S |
| Anti-MCU Rabbit mAb | 1:1000 | Cell Signaling Technology | 14997S |
| Anti-MICU1 (CBARA1) Rabbit mAb | 1:1000 | Custom-made antibody | Custom-made antibody |
| Anti-MICU2 Rabbit pAb | 1:500 | Bethyl Laboratories Inc | AB300-BL19212 |
| Anti-MICU3 Rabbit pAb | 1:1000 | Sigma Aldrich | HPA024771 |
| Anti-MCUB (CCDC109B) Rabbit pAb | 1:500 | Proteintech | 20387-1-AP |
| Anti-EMRE Rabbit pAb | 1:500 | Bethyl Laboratories Inc | AB300-BL19208 |
| Anti-MCUR1 Rabbit pAb | 1:500 | Invitrogen | PA5-32164 |
| Anti SLC24A6 (NCLX) Rabbit pAb | 1:1000 | Proteintech | 21430-1-AP |
| Anti-ATP5A1 Mouse mAb | 1:1000 | Proteintech | 66037-1-Ig |
| Anti-FLAG M2 mouse mAb | 1:1000 | Sigma-Aldrich | F1804 |
| Anti-rabbit IRDye 680 RD | 1:1000 | LICORbio | 926-68071 |
| Anti-rabbit IRDye 800 RD | 1:1000 | LICORbio | 926-32211 |
| Anti-mouse IRDye 680 RD | 1:1000 | LICORbio | 926-68070 |
| Anti-mouse IRDye 800 RD | 1:1000 | LICORbio | 926-32210 |

134

135 **Supplementary Table 7. Primary and secondary antibodies used in this study.** This table
136 summarizes antibody targets, host species, working dilutions, suppliers, and catalog numbers
137 for all primary and secondary antibodies used throughout the study.
